# Promoter architecture and developmental-expression divergence in the *Daphnia pulex* complex

**DOI:** 10.64898/2026.08.31.748272

**Authors:** Ruyun Deng, Feng Guo, Sarah Walls, Michael Lynch, Zhiqiang Ye

## Abstract

Changes in gene regulation are central to developmental evolution, yet how transcription-initiation architecture contributes to developmental expression divergence remains poorly understood. Here, we integrated chromosome-level genome assemblies, developmental RNA-seq, and STRIPE-seq-based transcription start site (TSS) profiling to examine regulatory divergence between two lineages of the *Daphnia pulex* complex, KAP4 and CON21, across embryonic, juvenile, and adult stages. The two lineages shared a broadly conserved developmental transcriptional program, but KAP4 consistently expressed more protein-coding genes and long non-coding RNAs than CON21. Genes with stage-specific expression were weakly expressed and showed elevated nucleotide diversity and sequence divergence relative to persistently expressed genes, consistent with relaxed purifying selection. Comparative transcriptomic analyses revealed strong stage-dependent expression divergence between the two lineages: the greatest similarity occurred during late embryonic to early juvenile development rather than mid-embryogenesis, providing limited support for a canonical developmental hourglass pattern. STRIPE-seq uncovered a conserved two-peak TSS organization upstream of annotated translation-start sites, but the relative use of distal and proximal initiation regions differed between lineages. Motif analyses further indicated that these initiation regions are associated with distinct *cis*-regulatory sequence environments. Promoter shape was closely linked to transcriptional output, with broad promoters showing the highest expression, peaked promoters the lowest, and intermediate promoters occupying an intermediate state. Together, these findings suggest that developmental regulatory divergence in the *D. pulex* complex is shaped by fine-scale remodeling of TSS usage and promoter architecture.

## Introduction

Developmental gene regulation is a fundamental mechanism shaping phenotypic diversity and evolutionary innovation. The development of multicellular organisms depends on coordinated spatial and temporal gene-expression programs that integrate maternally inherited factors, zygotic transcription, cellular differentiation, morphogenesis, and tissue specialization into complex developmental trajectories. Because developmental processes are largely controlled by when, where, and to what extent genes are expressed, evolutionary changes in gene regulation have long been recognized as a major source of developmental and phenotypic diversification (Raff 1996; Gerhart and Kirschner 1997; Carroll 2005). Comparative studies further show that conserved developmental or phenotypic outcomes can be maintained despite substantial divergence in the underlying genetic and regulatory mechanisms, a phenomenon commonly referred to as developmental system drift (True and Haag 2001). Consistent with this view, comparative transcriptomic studies across related species have shown that broadly conserved developmental programs can nevertheless accumulate lineage-specific changes in gene expression (Kalinka et al. 2010; Gerstein et al. 2014; Wang et al. 2020). Understanding how such regulatory divergence accumulates while developmental programs remain robust therefore remains a central question in evolutionary developmental biology.

A central construct addressing this issue is the developmental hourglass model, which proposes that early and late developmental stages are relatively divergent, whereas mid-embryonic stages are more conserved because they correspond to a phylotypic period characterized by stronger developmental constraints (Duboule 1994; Irie and Kuratani 2014). Comparative analyses of developmental transcriptomes have provided some support for this model, showing reduced gene-expression divergence during mid-embryogenesis among related *Drosophila* species (Kalinka et al. 2010). Moreover, evolutionary analyses suggest that developmental expression patterns are associated with patterns of sequence variation and divergence, consistent with stronger constraint acting on genes expressed during conserved developmental periods (Cruickshank and Wade 2008). However, the position and strength of the hourglass-like conservation pattern are not universal and may vary among taxa, developmental systems, and molecular layers. For example, hourglass-like temporal patterns of developmental conservation have also been observed at the level of microRNA expression (Ninova et al. 2014), whereas comparative studies across a wider range of taxa have revealed shifts in the timing of maximal conservation or deviations from the canonical mid-developmental conservation pattern (Drost et al. 2017). Together, these studies indicate that developmental conservation does not simply reflect uniform constraint, but instead emerges from dynamic interactions between conserved developmental requirements and lineage-specific regulatory remodeling.

Transcriptional regulation provides a mechanistic link between developmental constraint and evolutionary innovation. Transcription initiation represents one of the earliest and most important regulatory steps in gene expression. In eukaryotes, transcription begins at transcription start sites (TSSs), which are often clustered into transcription start regions (TSRs). These regions integrate core promoter elements, local sequence composition, chromatin accessibility, and transcription-factor binding to determine transcriptional output. Promoters can differ substantially in their initiation patterns. Some genes initiate transcription from narrow regions dominated by one or a few major TSSs, producing peaked promoters, whereas others initiate transcription across broader regions with more dispersed TSS usage, producing broad promoters (Carninci et al. 2006; Hoskins et al. 2011; Raborn et al. 2016; Policastro et al. 2020). These promoter architectures are associated with distinct regulatory properties, with broad promoters often linked to constitutive expression and peaked promoters frequently associated with regulated, tissue-specific, or developmental expression programs (Hoskins et al. 2011; Raborn et al. 2016; Haberle and Stark 2018).

Evolutionary changes in promoter architecture may therefore provide an important mechanism for modifying developmental expression programs without extensive changes to protein-coding sequences. Such changes can involve shifts in TSS usage, promoter shape, initiation strength, or regulatory motif composition. Although core promoter functions are often conserved across animals, promoter sequences and regulatory organization can evolve rapidly, providing a potential route for lineage-specific modulation of transcriptional output (Ohler 2006; Haberle et al. 2014; Haberle and Stark 2018). Recent advances in transcription-initiation profiling have enabled genome-wide characterization of TSS organization and promoter architecture across developmental contexts (Carninci et al. 2006; Cvetesic et al. 2018; Policastro et al. 2020). These approaches have expanded our ability to move beyond gene-level expression measurements and investigate how transcription initiation landscapes contribute to regulatory variation. However, most studies of promoter architecture and its evolution have focused on classical model organisms, particularly *Drosophila melanogaster* and mammals (Roy et al. 2010; Tomancak et al. 2007; Hoskins et al. 2011; Graveley et al. 2011; Lu et al. 2020), and how promoter architecture evolves across developmental trajectories in closely related lineages remains poorly understood. A major challenge is to determine how lineage-specific changes in transcription initiation architecture contribute to developmental expression divergence. Although comparative RNA-seq studies can identify genes and developmental stages with divergent expression patterns, they cannot distinguish whether these differences arise from altered TSS usage, promoter shape, regulatory motif turnover, or other cis-regulatory changes. Conversely, promoter profiling alone provides limited insight into how architectural differences influence developmental transcriptional output. Integrating developmental RNA-seq with genome-wide TSS profiling therefore provides a powerful framework for linking regulatory architecture with lineage-specific expression divergence.

The freshwater microcrustacean *Daphnia pulex* provides an excellent system for investigating the evolutionary basis of developmental regulatory divergence. *Daphnia* has long served as a model for ecology, evolution, toxicology, and developmental plasticity because of its ecological importance, phenotypic plasticity, and capacity for clonal reproduction under laboratory conditions (Ebert 2005; Colbourne et al. 2011; Lampert 2011). Extensive genomic resources, including high-quality genome assemblies, developmental datasets, and molecular tools, have further established *Daphnia* as an emerging system for studying evolutionary genomics and gene regulation (Eriksson et al. 2013; Naraki et al. 2013; Ye et al. 2017; Ye et al. 2019). The *D. pulex* species complex is particularly valuable because it contains morphologically similar but genetically divergent lineages, providing an opportunity to examine regulatory evolution in the context of largely conserved developmental programs. Although previous studies have characterized promoter architecture and adult sex-specific expression in *D. pulex* (Raborn et al. 2016), how promoter organization evolves across developmental trajectories and contributes to lineage-specific expression divergence remains unknown.

Here, we combine chromosome-level genome assemblies, RNA-seq, and STRIPE-seq– based TSS profiling to examine gene expression and promoter architecture during development in two lineages of the *D. pulex* complex: KAP4 from North America and CON21 from Europe. Across ten developmental stages spanning embryogenesis to adulthood, we investigate three major questions: 1) how developmental-expression programs diverge between lineages and whether they follow a canonical hourglass pattern; 2) whether stage-specific genes exhibit distinct evolutionary constraints; and 3) whether expression divergence is associated with lineage-specific promoter architecture and TSS organization. By integrating comparative transcriptomics with genome-wide transcription-initiation profiling, this study provides insights into how regulatory architecture contributes to developmental expression divergence between closely related lineages.

## Results

### Developmental transcriptome dynamics and altered selection on stage-specific genes

In this study, we analyzed two *Daphnia pulex* genomes representing distinct evolutionary lineages: KAP4, collected from Indiana, USA, and CON21, collected from Europe. Both genomes were assembled to the chromosome level, with estimated genome sizes of 133.2 Mb and 143.8 Mb, respectively, and each assembly contained 12 chromosomes (**Table S1**). BUSCO analyses (Simão et al. 2015) indicated high completeness for both assemblies, with scores of 98.9% for KAP4 and 96.6% for CON21. Genome annotation identified 15,282 protein-coding genes (PCGs) and 4,132 long non-coding RNAs (lncRNAs) in KAP4, compared with 14,492 PCGs and 2,548 lncRNAs in CON21 (**Table S1**). Thus, despite its larger genome size, CON21 contained fewer annotated protein-coding genes and lncRNAs than KAP4, reflecting differences in annotated gene content between the two assemblies. A total of 11,695 orthologous gene pairs were identified between KAP4 and CON21. Sequence divergence analysis of these orthologs revealed mean nonsynonymous (*K*a) and synonymous (*K*s) substitution rates of 0.0081 and 0.0474, respectively.

We next quantified gene-expression dynamics in the two lineages across ten developmental stages, spanning embryogenesis (A–F), the juvenile period (G–I), and adulthood (M) (**Figure 1A**). Embryogenesis was divided into six consecutive stages defined by distinct morphological features (see Methods). Following release from the brood chamber, juveniles were collected at three time points: stage G at 8 h (approximately instar 1), stage H at 56 h (approximately instar 2), and stage I at 104 h (approximately instar 4). Adults (stage M) were defined as individuals maintained for seven days after release, at which point ovaries are well developed. For each lineage, RNA-seq was performed with three independent biological replicates (**Figure 1B**), generating an average of 76 million mapped reads per sample, providing high sequencing depth for transcript quantification (Conesa et al. 2016). Biological replicates showed high reproducibility, with mean Pearson correlation coefficients of 0.96 for both KAP4 and CON21 (**Table S2**).

**Figure 1.**
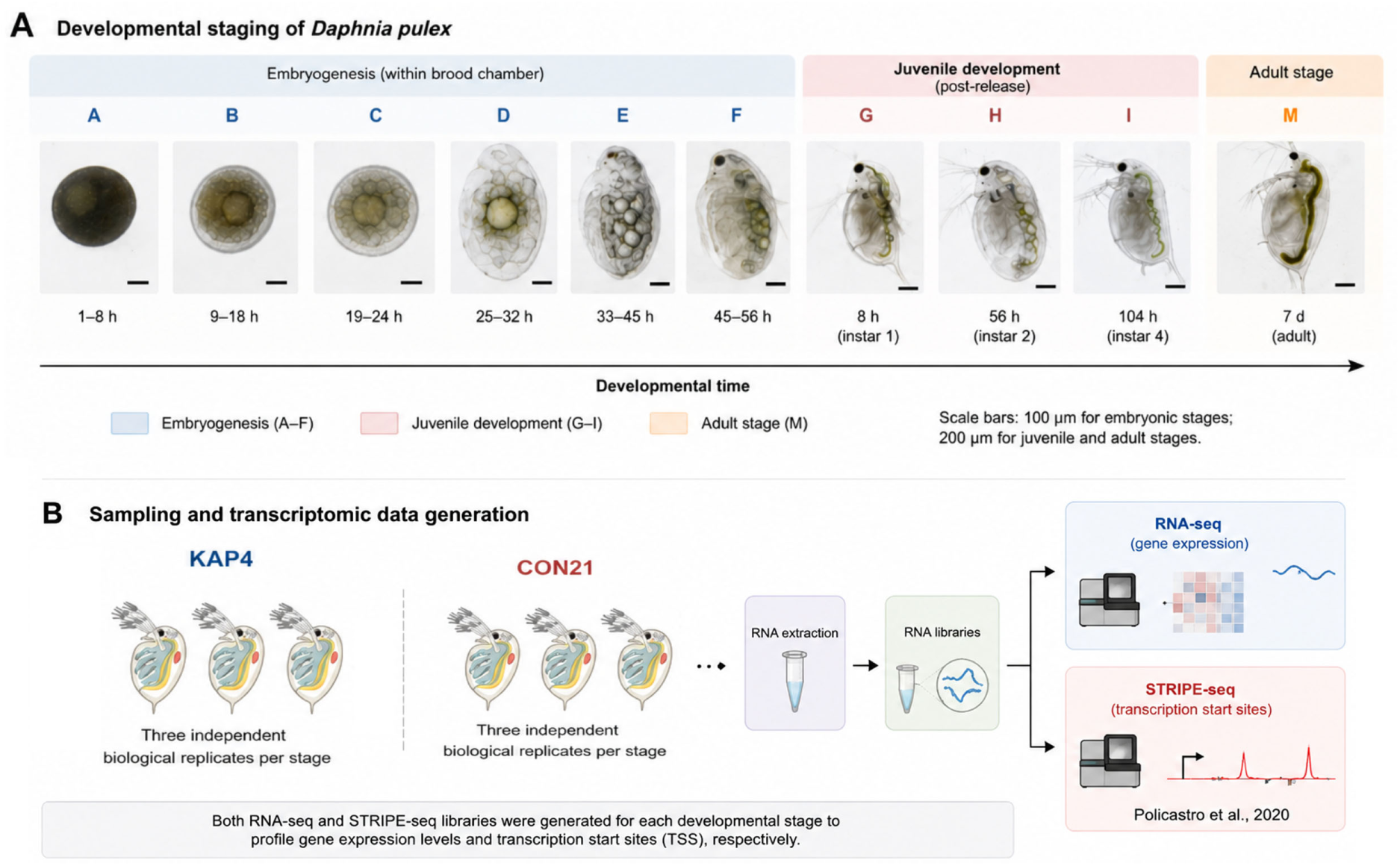
Developmental staging and transcriptomic sampling design. **A)** Overview of the ten developmental stages analyzed in this study, spanning embryogenesis (stages A–F), juvenile development (stages G–I), and adulthood (stage M). Embryonic stages were defined based on morphological criteria, whereas juvenile and adult stages were defined by time after release from the brood chamber. **B)** Summary of RNA-seq and STRIPE-seq sampling for the two *Daphnia pulex* lineages, KAP4 and CON21. RNA-seq libraries were generated from three independent biological replicates for each developmental stage. STRIPE-seq libraries were generated from matched developmental samples where available and used to profile transcription start sites.

Using transcripts per million (TPM) > 1 as the expression threshold for PCGs, we detected 13,652 expressed PCGs in KAP4 (89% of all annotated genes) and 12,871 in CON21 (89%) (**Supplementary File**). Thus, although KAP4 contains more annotated PCGs, the overall fraction of the PCG repertoire expressed during development was nearly identical between the two lineages. For lncRNAs, using a lower threshold of TPM > 0.2 to account for their generally lower expression, 3,584 lncRNAs (87% of annotated lncRNAs) were detected in KAP4 and 2,203 (86%) in CON21.

Across individual developmental stages, both lineages showed a general increase in the number and proportion of expressed PCGs from early embryogenesis toward later development. In KAP4, 9,108–10,031 PCGs were expressed during stages A–C, corresponding to approximately 60–66% of the annotated PCG repertoire, whereas more than 12,000 genes (>79%) were expressed at several later stages, including F, H, and M (**Table 1**). CON21 showed a broadly similar developmental trend, with 7,836–8,320 PCGs expressed during stages A–C (54–57% of annotated PCGs). The proportion of expressed PCGs subsequently increased from 64.0% at stage D to a maximum of 82% at stage I, followed by a decline to 69% at stage M (**Table 1**). Although the absolute number of expressed PCGs was generally lower in CON21, this difference partly reflects the smaller number of annotated PCGs in that assembly. The similar developmental increase in the fraction of expressed genes therefore provides a more comparable measure of transcriptional dynamics between the two lineages. Consistent with this overall similarity, 76–85% of expressed PCGs were shared between KAP4 and CON21 at corresponding developmental stages (**Figure 2A**), indicating substantial conservation of the active protein-coding transcriptome.

**Figure 2.**
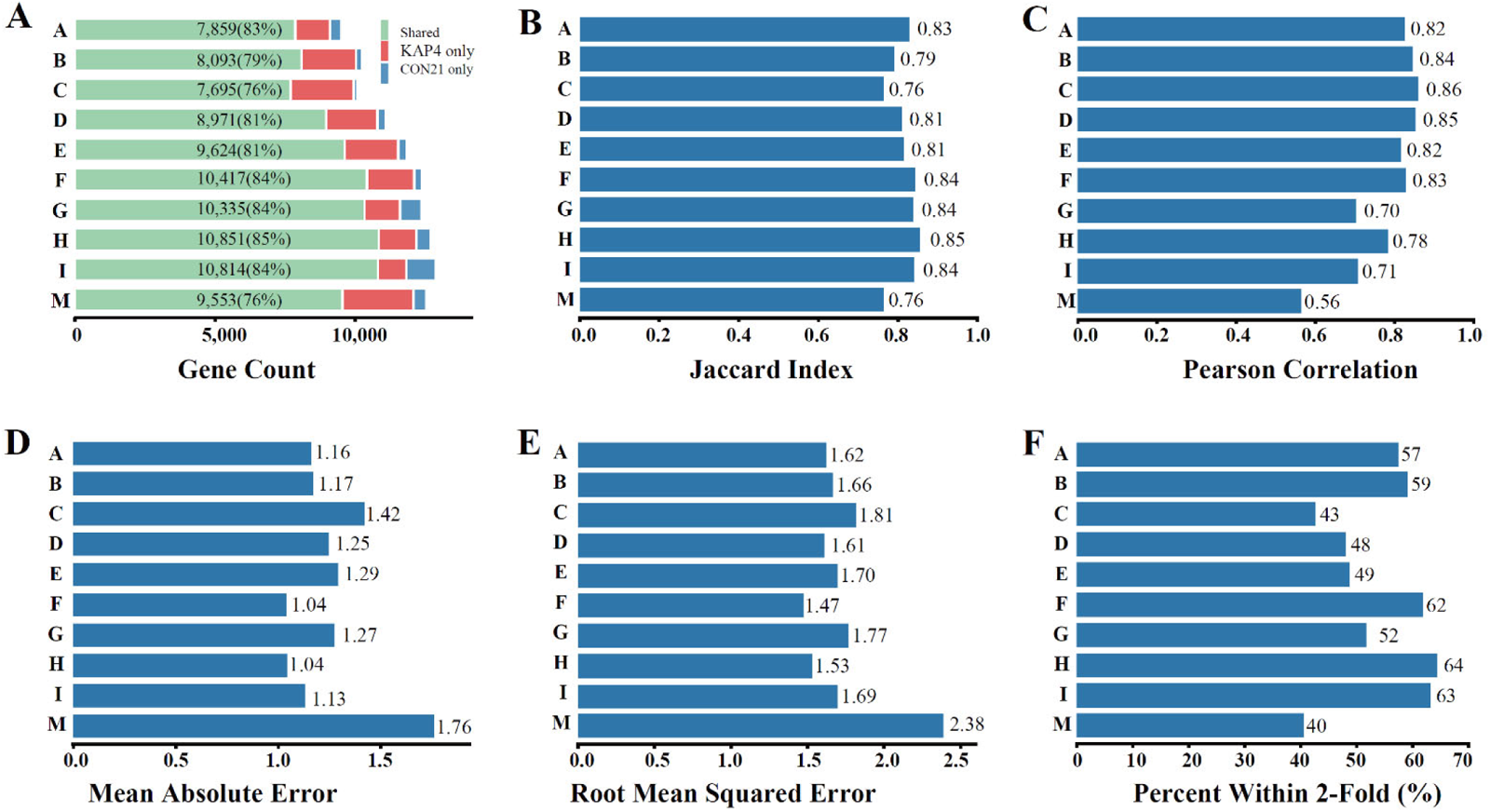
Stage-dependent developmental transcriptome divergence between KAP4 and CON21. **(A)** Numbers of expressed genes shared between KAP4 and CON21 or expressed only in one lineage at each developmental stage. **(B)** Jaccard index measuring proportional overlap of expressed gene sets between KAP4 and CON21. **(C)** Pearson correlation coefficients for relative expression levels of shared expressed genes. **(D)** Mean absolute error (MAE) quantifying the average absolute difference in expression levels. **(E)** Root mean squared error (RMSE) highlighting larger deviations in expression magnitude. **(F)** Percentage of genes with expression differences within twofold between KAP4 and CON21.

**Table 1.** Summary of expressed protein-coding genes (PCGs) and long non-coding RNAs (lncRNAs) across developmental stages in KAP4 and CON21. Percentages indicate the proportion of expressed genes relative to the total number of annotated PCGs or lncRNAs in each lineage. “Specific” denotes genes expressed exclusively at a single developmental stage. Mean TPM values represent the expression levels of expressed PCGs and lncRNAs across developmental stages. The TPM ratio was calculated as the mean TPM of expressed PCGs divided by that of expressed lncRNAs.

| Stage | Expressed PCG | TPM PCG | #Specific PCG | Expressed lncRNA | TPM lncRNA | #Specific lncRNA | TPM Ratio |
| --- | --- | --- | --- | --- | --- | --- | --- |
| **KAP4** |  |  |  |  |  |  |  |
| A | 9,108 (60%) | 106.2 | 52 | 2,868 (69%) | 8.8 | 34 | 12.1 |
| B | 10,031 (66%) | 96.4 | 37 | 2,779 (67%) | 10.2 | 3 | 9.5 |
| C | 9,946 (65%) | 97.8 | 14 | 2,678 (65%) | 8.2 | 9 | 11.9 |
| D | 10,800 (71%) | 89.6 | 18 | 2,906 (70%) | 8.3 | 9 | 10.8 |
| E | 11,532 (75%) | 84.2 | 30 | 2,596 (63%) | 9.2 | 1 | 9.2 |
| F | 12,106 (79%) | 79.1 | 57 | 2,933 (71%) | 11.9 | 3 | 6.6 |
| G | 11,603 (76%) | 82.8 | 44 | 2,465 (60%) | 15.0 | 4 | 5.5 |
| H | 12,180 (80%) | 78.6 | 27 | 2,874 (70%) | 12.6 | 8 | 6.2 |
| I | 11,829 (77%) | 80.8 | 15 | 2,678 (65%) | 13.1 | 1 | 6.2 |
| M | 12,079 (79%) | 78.6 | 129 | 3,225 (78%) | 11.8 | 55 | 6.7 |
| mean | 11,121 (73%) | 87.4 | 42 | 2,800 (68%) | 10.9 | 13 | 8.5 |
| **CON21** |  |  |  |  |  |  |  |
| A | 8,261 (57%) | 117.9 | 104 | 1,270 (50%) | 18.7 | 136 | 6.3 |
| B | 8,320 (57%) | 116.2 | 19 | 912 (36%) | 33.7 | 6 | 3.4 |
| C | 7,836 (54%) | 122.2 | 1 | 855 (34%) | 43.2 | 9 | 2.8 |
| D | 9,274 (64%) | 103.7 | 13 | 918 (36%) | 37.2 | 5 | 2.8 |
| E | 9,939 (69%) | 96.6 | 46 | 1,083 (43%) | 31.0 | 13 | 3.1 |
| F | 10,697 (74%) | 88.5 | 31 | 1,215 (48%) | 33.1 | 9 | 2.7 |
| G | 11,098 (77%) | 85.0 | 79 | 1,427 (56%) | 27.0 | 36 | 3.1 |
| H | 11,372 (78%) | 83.4 | 22 | 1,402 (55%) | 21.7 | 7 | 3.8 |
| I | 11,869 (82%) | 79.7 | 107 | 1,683 (66%) | 16.6 | 89 | 4.8 |
| M | 10,009 (69%) | 94.3 | 104 | 1,090 (43%) | 47.1 | 66 | 2.0 |
| mean | 9,868 (68%) | 98.7 | 53 | 1,186 (47%) | 30.9 | 38 | 3.5 |

lncRNAs exhibited a developmental pattern distinct from that of PCGs. Across developmental stages, KAP4 consistently expressed more lncRNAs than CON21, with an average of 2,800 expressed lncRNAs per stage compared with 1,186 in CON21 (**Table 1**). When normalized by the total number of annotated lncRNAs in each lineage, KAP4 expressed an average of 68% of its lncRNA repertoire per stage, whereas CON21 expressed 47%. In KAP4, the proportion of expressed lncRNAs remained relatively stable across development, ranging from 60% to 78% (2,465–3,225 lncRNAs), with the highest proportion observed in adults (stage M). By contrast, CON21 showed greater stage-to-stage variation, with 34–66% of annotated lncRNAs expressed (855–1,683 lncRNAs), reaching the highest proportion at stage I (**Table 1; Figure S1**). Differences between the two lineages were most pronounced during early and middle development, when KAP4 expressed both a larger absolute number and a greater proportion of its annotated lncRNA repertoire. KAP4 also contained a larger fraction of lineage-specific expressed lncRNAs across stages (20–54%; **Figure S2**). Because the two genomes differ substantially in their total numbers of annotated lncRNAs, differences in absolute counts should be interpreted cautiously. Nevertheless, the consistently higher proportion of expressed lncRNAs in KAP4, together with the greater stage-to-stage variation observed in CON21, indicates stronger lineage-specific divergence in developmental lncRNA expression than was observed for PCGs.

In addition to differences in the number of expressed transcripts, PCGs and lncRNAs also differed markedly in expression intensity. In KAP4, the mean expression level of PCGs was approximately 8.5-fold higher than that of lncRNAs (**Table 1**). Combined with the larger number of expressed PCGs, this resulted in an estimated 34-fold greater aggregate PCG expression than lncRNA expression. A similar pattern was observed in CON21, where aggregate PCG expression was approximately 30-fold greater than that of lncRNAs. Thus, although lncRNAs showed stronger lineage- and stage-specific variation in transcript representation, they contributed a much smaller fraction of the overall transcriptional output than PCGs in both lineages.

We next classified PCGs into persistently expressed and stage-specific categories, allowing us to distinguish broadly active genes from genes potentially associated with specific developmental transitions. A total of 7,649 genes (56% of all expressed genes) in KAP4 and 6,842 genes (53%) in CON21 were expressed across all developmental stages (**Figure S1**). Functional-enrichment analyses showed that these persistently expressed PCGs had broadly similar functional profiles in the two lineages, with enrichment for GO categories related to translation and ribosomal structure, RNA binding, cytosolic and mitochondrial functions, membrane components, extracellular region or matrix organization, proteolysis, serine-type endopeptidase activity, and cuticle-related structural components (**Figures S3 and S4**). These shared functional profiles are consistent with persistent genes contributing predominantly to fundamental cellular and developmental processes required across multiple developmental stages.

Stage-specific PCGs, in contrast, were relatively uncommon and displayed distinct temporal distributions across development. In KAP4, the largest number of stage-specific PCGs was observed at stage M (129 genes), whereas CON21 showed the highest number at stage I (107 genes), followed closely by stage M (104 genes) (**Table 1**). These genes were also expressed at substantially lower levels than persistently expressed PCGs, with a mean TPM of approximately 11 across developmental stages compared with approximately 106 for persistent genes (**Table 2**). Stage-specific lncRNAs showed even greater lineage-specific differences. In KAP4, 1,682 lncRNAs (41%) were expressed across all stages, compared with 432 (17%) in CON21 (**Figure S1**), while the distribution of stage-specific lncRNAs varied markedly across development (**Table 1**). Overall, stage-specific expression tended to be more pronounced at the earliest developmental stage and in adults. Functional-enrichment analyses showed that stage-specific PCGs were associated with temporally restricted developmental and physiological processes. In both lineages, early-stage genes were enriched for ubiquitin/proteasome-related protein turnover, whereas stage E showed shared enrichment for glycosylation-related processes. Later-stage genes were more frequently associated with membrane functions, extracellular structures, and stress responses (**Figures S5 and S6**). Lineage-specific differences were also evident: KAP4 showed enrichment for chromatin-associated processes at stage C and cuticle- and chitin-related extracellular matrix functions at stages F and M, whereas CON21 showed stronger enrichment for regulatory DNA binding, neuropeptide signaling, sodium-ion transport, and oxidative-stress responses (**Figures S5 and S6**). Together, these patterns contrast with the broadly conserved cellular functions of persistently expressed genes and indicate that stage-specific PCGs are more closely associated with temporally and lineage-restricted developmental functions.

**Table 2.** Average population-genetic parameter estimates for protein-coding genes expressed specifically in each developmental stage (A–I, M) in KAP4. TPM refers to transcripts per million. “# π” and “# d” denote the numbers of genes for which nucleotide diversity (π) and sequence divergence (d) were estimated. “Background genes” refer to genes with persistent expression, for which TPM values represent averages across all stages. The values in parentheses represent the standard errors (SE).

| KAP4 | TPM | # $\pi$ | $\pi_N$ | $\pi_S$ | $\pi_N/\pi_S$ | # $d$ | $d_N$ | $d_S$ | $d_N/d_S$ |
| --- | --- | --- | --- | --- | --- | --- | --- | --- | --- |
| A | 41.9 (18.7) | 34 | 0.005 (0.001) | 0.020 (0.002) | 0.545 (0.213) | 28 | 0.039 (0.003) | 0.160 (0.011) | 0.287 (0.050) |
| B | 17.5 (7.0) | 32 | 0.004 (0.001) | 0.017 (0.002) | 0.885 (0.355) | 21 | 0.030 (0.004) | 0.152 (0.012) | 0.230 (0.038) |
| C | 2.7 (0.6) | 5 | 0.005 (0.001) | 0.015 (0.004) | 1.237 (0.906) | 2 | 0.046 (0.015) | 0.243 (0.072) | 0.187 (0.007) |
| D | 2.2 (0.3) | 7 | 0.007 (0.003) | 0.028 (0.005) | 0.276 (0.120) | 5 | 0.022 (0.006) | 0.176 (0.020) | 0.138 (0.041) |
| E | 5.2 (1.7) | 24 | 0.007 (0.001) | 0.039 (0.005) | 0.371 (0.125) | 16 | 0.028 (0.004) | 0.183 (0.012) | 0.180 (0.048) |
| F | 3.8 (0.6) | 43 | 0.006 (0.001) | 0.021 (0.002) | 0.588 (0.159) | 32 | 0.035 (0.004) | 0.150 (0.009) | 0.257 (0.029) |
| G | 11.5 (3.1) | 33 | 0.005 (0.001) | 0.026 (0.002) | 0.228 (0.031) | 27 | 0.031 (0.003) | 0.170 (0.010) | 0.195 (0.029) |
| H | 2.4 (0.3) | 17 | 0.005 (0.001) | 0.023 (0.003) | 0.278 (0.046) | 12 | 0.032 (0.004) | 0.122 (0.007) | 0.270 (0.032) |
| I | 3.0 (0.8) | 12 | 0.005 (0.001) | 0.024 (0.004) | 0.338 (0.104) | 9 | 0.035 (0.007) | 0.208 (0.033) | 0.192 (0.044) |
| M | 20.9 (6.8) | 95 | 0.006 (0.000) | 0.022 (0.002) | 0.451 (0.073) | 66 | 0.039 (0.002) | 0.160 (0.006) | 0.274 (0.023) |
| **mean** | **11.1** | **30** | **0.006** | **0.024** | **0.520** | **22** | **0.034** | **0.172** | **0.221** |
| Bkgd | 105.6 | 7,328 | 0.002 | 0.014 | 0.246 | 6,889 | 0.016 | 0.122 | 0.120 |

Finally, we examined whether stage-specific PCGs experience different evolutionary constraints from persistently expressed genes. To assess variation at both population and interspecific timescales, we analyzed nucleotide diversity within *D. pulex* and sequence divergence from *D. obtusa* orthologs, with an average silent-site divergence of approximately 0.13 between the two species. Extensive population-genomic data are available for North American *D. pulex*, comprising approximately 1,800 clones (Ye et al. 2023; Lynch et al. 2024). Because comparable population-level data are not available for CON21, polymorphism-based analyses were restricted to KAP4. Stage-specific genes were compared with genes expressed consistently across all developmental stages, hereafter referred to as background genes. Stage-specific genes show substantially elevated nucleotide diversity for both synonymous and nonsynonymous sites (**Table 2**). In KAP4, the mean π_N_ and π_S_ of stage-specific genes across stages are 0.006 and 0.024, respectively, compared with 0.002 and 0.014 for background genes (**Table 2**). This results in an approximately twofold elevation in π_N_/π_S_ in stage-specific genes relative to the background level, which suggests reduced efficiency of purifying selection on nonsynonymous variation under the assumption that silent sites are behaving in a neutral fashion. However, the reduced π_S_ for background genes suggests that even silent sites may be under purifying selection, which would imply an even greater disparity than expected under the assumption of neutrality. Stage-specific genes also showed consistently elevated sequence divergence. Across stages, stage-specific genes had approximately 1.8-fold higher d_N_/d_S_ ratios than background genes (**Table 2**), further indicating relaxed selective constraint over evolutionary time. Together, these results suggest that stage-specific genes tend to be weakly expressed and evolve under relaxed purifying selection, as reflected by elevated polymorphism and divergence, particularly at nonsynonymous sites.

### Stage-dependent expression divergence between KAP4 and CON21

To assess how developmental-expression programs diverge between KAP4 and CON21, we compared gene-expression profiles across ten developmental stages using multiple complementary metrics. These metrics were designed to capture different dimensions of transcriptomic similarity, including overlap in expressed-gene repertoires, conservation of relative expression patterns, and absolute differences in expression levels.

First, we quantified the number of genes expressed in both KAP4 and CON21 at each corresponding developmental stage. This shared expressed gene count reflects the size of the intersection between the two sets of expressed genes and therefore measures how many genes are transcriptionally active in both taxa at the same stage. Across all stages, the largest number of shared expressed genes was observed at stage H, with 10,851 genes expressed in both KAP4 and CON21 (**Figure 2A**). Stages F, G, and I also showed high numbers of shared expressed genes, indicating broad conservation of active gene repertoires during late embryonic and juvenile development. In contrast, stage C had the fewest shared expressed genes, with 7,695 genes detected in both lineages (**Figure 2A**). Thus, based on shared gene counts alone, similarity in the active transcriptome was highest during late embryonic to juvenile stages and lowest during stage C.

Because the total number of expressed genes varies among developmental stages and between lineages, we also calculated the Jaccard index, defined as the number of genes expressed in both lineages divided by the total number of genes expressed in at least one lineage. This is a measure of the proportional overlap of expressed-gene sets, with higher values indicating that the two taxa activate a more similar set of genes at a given developmental stage. Consistent with the shared-gene count analysis, stage H showed the highest Jaccard index (0.85; **Figure 2B**), indicating the greatest proportional overlap in expressed-gene repertoires. Stages F, G, and I also shared nearly equal high Jaccard indices of 0.84. In contrast, stages M and C showed the lowest Jaccard index (0.76). These results indicate that expressed-gene composition was most similar between KAP4 and CON21 during late embryonic to juvenile development, whereas adulthood and stage C showed reduced overlap in active gene repertoires.

We next compared relative expression structure using Pearson correlation based on log_2_(TPM + 1)-transformed expression values. Unlike shared-gene counts and Jaccard indices, which focus on whether genes are expressed or not, Pearson correlations were calculated using orthologous genes expressed in both lineages at each developmental stage and thus measure conservation of relative expression patterns among shared genes. The highest Pearson correlation was observed at stage C (0.86), followed by stage D (0.85) and stage B (0.84) (**Figure 2C**). These results indicate that early to mid-embryonic stages retained relatively conserved expression structure among shared expressed genes. However, this high correlation does not necessarily indicate global transcriptomic conservation, because stage C also had low expressed-gene overlap and large absolute expression differences. In contrast, stage M showed the lowest Pearson correlation (0.56), indicating that adult expression profiles were the most divergent in terms of relative expression structure.

To quantify absolute expression divergence, we calculated mean absolute error (MAE) and root mean squared error (RMSE). Stage F showed the lowest MAE (1.04), with stage H showing a similarly low value (1.04), whereas stage M had the highest MAE (1.76) and stage C the highest among embryonic stages (1.42) (**Figure 2D**). RMSE produced the same qualitative pattern (**Figure 2E**). As a complementary and more intuitive, threshold-based measure of similarity, we also calculated the proportion of genes with expression differences within twofold between KAP4 and CON21. Stage H had the highest proportion of genes within twofold expression difference (64%), followed by stage I (63%) and stage F (62%), whereas stage M had the lowest proportion (40%), followed by stage C (43%) (**Figure 2F**). These results further support the conclusion that late embryonic to juvenile stages, particularly stages H and F, show the closest gene-level expression similarity, whereas stages M and C show stronger expression divergence.

Overall, the different metrics captured complementary dimensions of transcriptomic similarity. Stage H showed the greatest overlap in expressed-gene repertoires, whereas stages F and H showed the smallest absolute expression differences. In contrast, stage M showed the strongest divergence across most metrics. Stage C represented a notable exception: although expression rankings were highly correlated between KAP4 and CON21, this stage showed relatively low expressed-gene overlap and substantial differences in expression magnitude. Thus, the greatest overall transcriptomic similarity between KAP4 and CON21 occurred during late embryonic to juvenile development, particularly at stages F and H, rather than during mid-embryogenesis.

### Lineage-specific differences in transcription initiation and promoter architecture

To investigate the potential regulatory basis of developmental expression differences, we generated TSS data using STRIPE-seq (Policastro et al. 2020). As in RNA-seq analysis, we quantified transcription start sites in two lineages, KAP4 and CON21, across ten developmental stages, including embryonic stages (A–F), juvenile stages (G–I), and the adult stage (M). Sequencing yielded sufficient coverage for genome-wide TSS profiling in both lineages, with an average of approximately 60 million reads per sample. Due to unsuccessful library construction, stage G was missing from KAP4 and stage M was missing from CON21. These missing stages were excluded from direct stage-to-stage comparisons. Because promoter usage is less well characterized for lncRNAs, subsequent analyses were restricted to protein-coding genes.

In KAP4, an average of 8,397 genes per stage were associated with detected TSSs, corresponding to 76% of expressed genes across stages. In CON21, an average of 7,045 genes per stage had detected TSSs, representing 73% of expressed genes (**Table S3**). Both lineages thus showed broad TSS coverage across development, although KAP4 exhibited a larger number of genes with detected TSSs. The positional distribution of TSS signals relative to annotated translation-start sites showed both shared and clone-specific patterns (**Figure 3**). Across most developmental stages, TSS profiles in both KAP4 and CON21 exhibited a characteristic double-peak structure upstream of annotated translation start sites (**Figure 3**). One peak was located in the distal upstream region, approximately −120 to −70 bp from the annotated translation start site, whereas the second peak was located closer to the start codon, approximately −40 to −10 bp (**Figure 3**), although the exact peak positions varied slightly among stages. The two peaks were observed across the available stages in both lineages, although their relative heights varied across development. The primary difference between lineages was the relative usage of the two peaks: in CON21, the proximal peak closer to the start codon generally showed higher relative frequency, whereas in KAP4, the distal upstream peak was often more pronounced (**Figure 3**).

**Figure 3.**
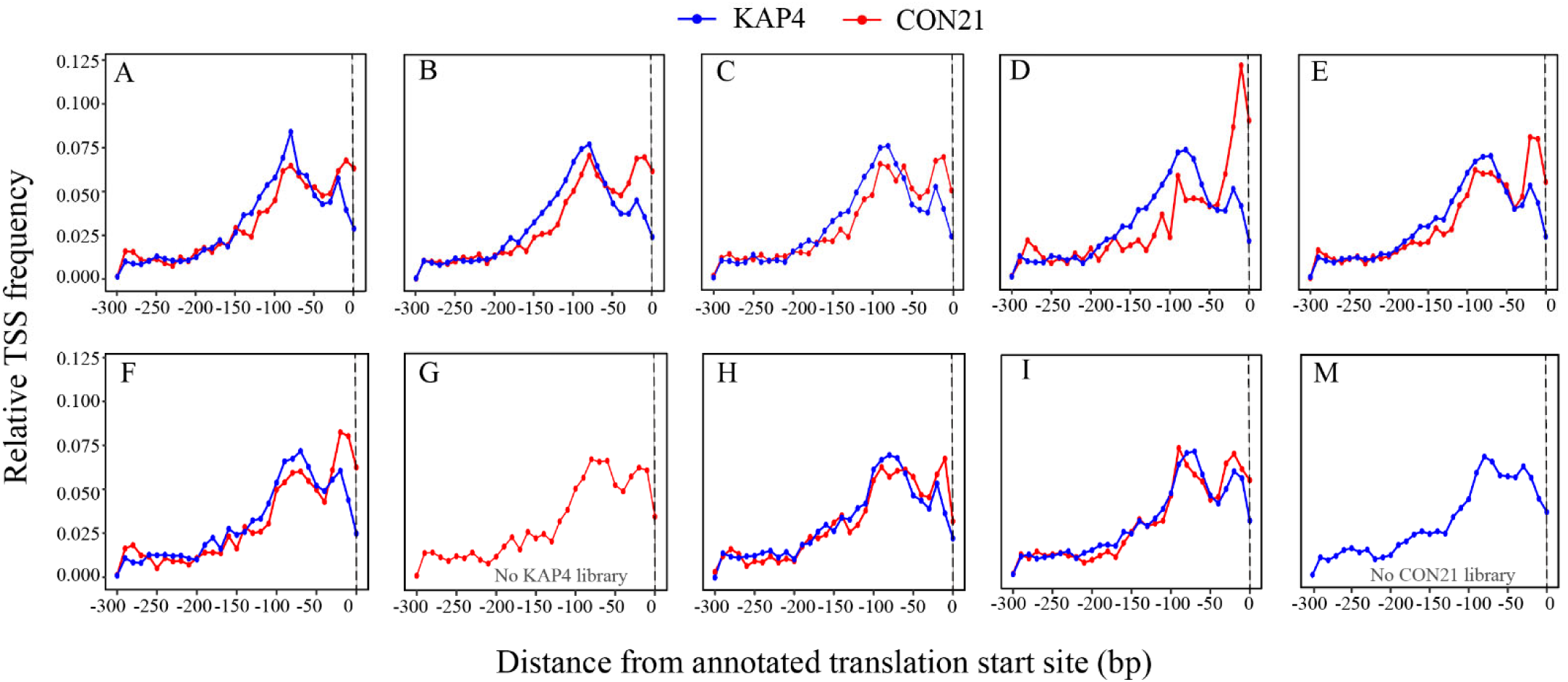
Stage-specific distributions of transcription start site (TSS) signals in KAP4 and CON21. Relative TSS frequencies were calculated within the 300 bp upstream region of annotated translation start sites. Blue lines represent KAP4 and red lines represent CON21. Each panel corresponds to a developmental stage (A–M). Stage G was unavailable for KAP4, and stage M was unavailable for CON21 because of unsuccessful library construction. The dashed vertical lines indicate the annotated translation-start site. Both lineages exhibited enrichment of TSS signals within proximal upstream regions, although KAP4 generally showed broader upstream distributions, whereas CON21 displayed more spatially confined initiation profiles in several developmental stages.

The relative prominence of the two peaks also changed across developmental stages, indicating stage-dependent modulation of transcription initiation. In early embryonic stages, the distal and proximal peaks were more comparable in height, whereas in later embryonic stages, especially D-F, the proximal peak became more prominent in CON21 (**Figure 3**). In juvenile stages, the two lineages showed more similar peak structures at stages H and I (**Figure 3**), although CON21 retained relatively strong proximal enrichment. Because stage G was unavailable for KAP4 and stage M was unavailable for CON21, direct lineage comparisons at these stages were not possible. Overall, these results suggest that KAP4 and CON21 share a conserved two-peak TSS distribution, but differ in the relative use of distal versus proximal initiation regions, with additional stage-dependent shifts in peak prominence across development.

Because aggregate TSS profiles could potentially be influenced by multiple initiation regions within the same gene, we next clustered neighboring CTSSs into transcription start regions (TSRs) and quantified the number of TSRs associated with each gene within each developmental stage (**see Methods**). On average, only 262 genes (3%) in KAP4 and 144 genes (2%) in CON21 were associated with two TSRs per stage (**Table S3**). Thus, genes containing two distinct TSRs represented only a small fraction of genes with detected transcription initiation, indicating that the aggregate bimodal distribution is unlikely to be driven primarily by individual genes carrying two separate initiation regions.

We next examined whether these differences in TSS usage were accompanied by changes in promoter architecture. For each TSR, promoter shape was quantified using the Shape Index (SI) and classified as Peaked, Intermediate, Broad, or Unclassified based on Gaussian mixture modeling (**see Methods**). Across the available developmental stages, promoter landscapes in both KAP4 and CON21 were dominated by Peaked promoters, whereas Broad promoters represented a relatively small fraction of total TSRs (**Figure 4**). Despite this shared overall pattern, the developmental distribution of promoter classes differed between the two lineages. In KAP4, the proportion of peaked promoters was lower during early stages, particularly stages B and C, where intermediate and unclassified promoters contributed a larger fraction (**Figure 4A**). As development progressed, peaked promoters became increasingly dominant, accompanied by a reduction in unclassified promoters. In contrast, CON21 showed a generally higher proportion of peaked promoters across most stages, especially at stages D, H, and I (**Figure 4B**). Although stages B and C also showed relatively lower peaked-promoter proportions in CON21, the contribution of unclassified promoters was less pronounced than in KAP4. Changes in the relative proportions of peaked and broad promoters across stages indicate developmental shifts in promoter-class composition. These shifts may reflect either stage-specific activation of genes with different promoter architectures or promoter-type switching within the same genes, which we examine further below. Together, these results indicate that KAP4 and CON21 share a broadly similar promoter-shape landscape dominated by peaked promoters, but differ in the developmental dynamics of promoter-class composition. KAP4 appears to undergo a more gradual developmental shift toward peaked promoter usage, whereas CON21 maintains a relatively high proportion of peaked promoters across most available stages. These patterns suggest lineage-associated differences in the developmental organization of promoter architecture.

**Figure 4.**
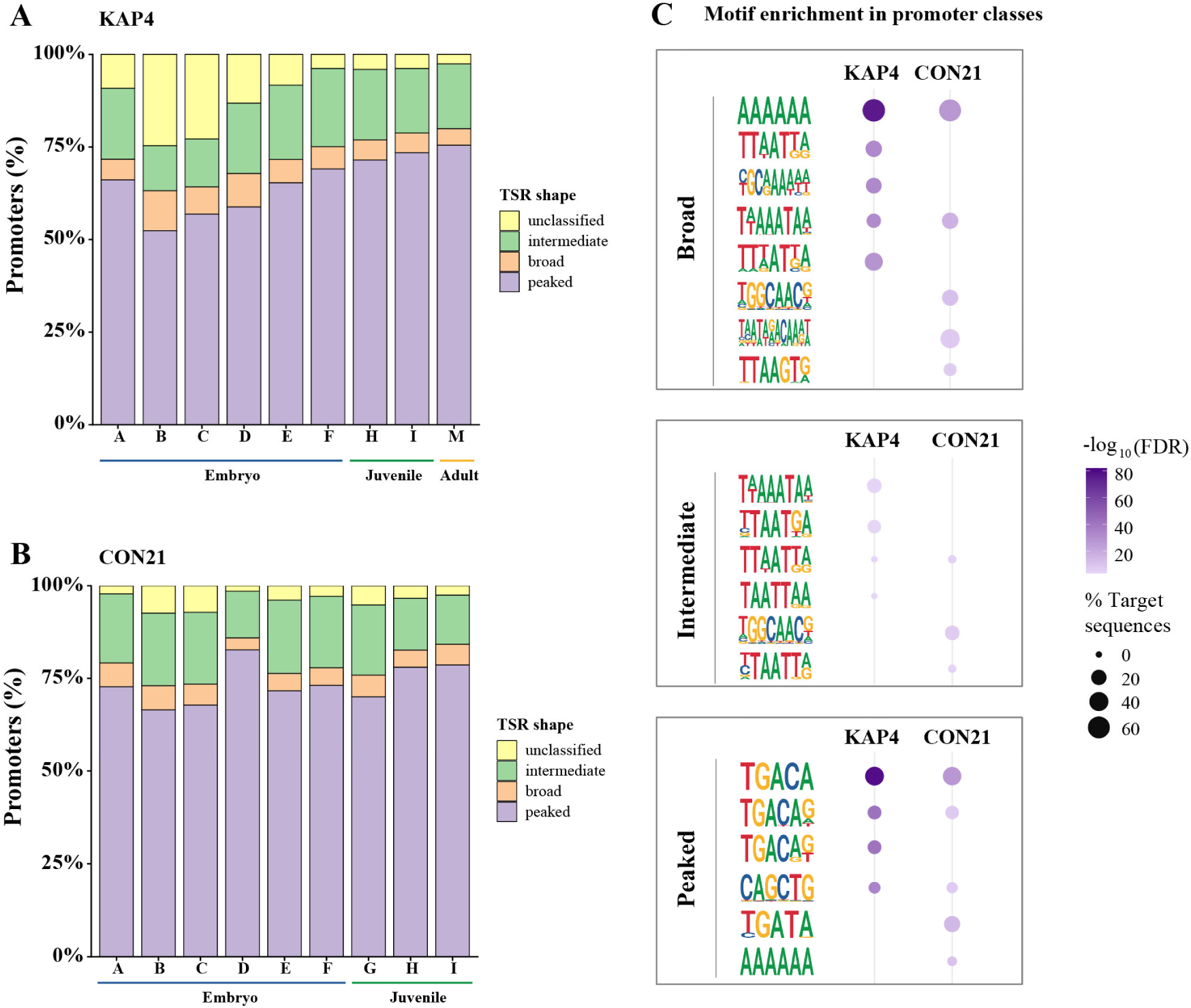
Developmental dynamics of promoter-shape classes and motif enrichment in KAP4 and CON21. **(A)** Proportional distribution of promoter-shape classes across developmental stages in KAP4. Stage G was unavailable for KAP4 and stage M was unavailable for CON21 due to unsuccessful library construction. **(B)** Corresponding promoter-shape distributions across developmental stages in CON21. Stages A–F correspond to embryonic development and stages G–I to juvenile development. Stage M was unavailable for CON21 due to unsuccessful library construction. **(C)** Enriched sequence motifs associated with different promoter classes in KAP4 and CON21. For each promoter class and lineage, the five most significantly enriched motifs are shown as sequence logos.

To explore the sequence features underlying promoter architecture, we compared cis-regulatory motifs among promoter-shape classes. Motif-enrichment analysis revealed promoter-class-specific patterns in both lineages (**Figure 4C**). Broad promoters were mainly associated with A/T-rich motifs, including poly-A-like and TAA/TTA-containing sequences, with AAAAAA-like motifs among the strongest signals. Peaked promoters showed a different motif profile, including motifs containing TGA- or TGACA-like sequence cores. Intermediate promoters showed mixed motif features, including both A/T-rich and TGA-related elements, consistent with their intermediate position in promoter-shape classification. Overall, motif-enrichment patterns differed more strongly among promoter-shape classes than between lineages, indicating that promoter shape is a major axis of *cis*-regulatory sequence differentiation in both KAP4 and CON21.

In a separate analysis from the promoter-shape motif comparison above, we asked whether the two peaks in the aggregate TSS distribution were associated with different local sequence environments. We therefore compared motif composition within the genomic intervals corresponding to the proximal (−40 to −10 bp) and distal (−120 to −70 bp) peaks in KAP4 and CON21 (**Figure S7**). These analyses were performed on sequences falling within the two positional intervals across genes, rather than by dividing genes into separate proximal- and distal-TSS groups. The two intervals showed clearly different motif-enrichment patterns. The two regions showed clearly different motif-enrichment patterns. In the proximal region, enriched motifs were dominated by A/T-rich sequences, including TAA/TTA-containing and other AT-rich motifs. Several of these motifs showed sequence characteristics reminiscent of TATA-like core-promoter elements, although they did not uniformly conform to the canonical TATA-box consensus. Consistent with this pattern, TATA-like sequences were more frequent in the proximal than in the distal region in both lineages, occurring in 10.78% versus 7.32% of CON21 sequences and 9.56% versus 6.30% of KAP4 sequences, respectively (**Table S4**).

In classical animal promoters, the TATA box is typically positioned upstream of the transcription start site and contributes to positioning the pre-initiation complex, whereas the initiator (Inr) element is associated more directly with the site of transcription initiation. We therefore also examined the occurrence of Inr-like sequences in the two initiation regions. Inr-like motifs were relatively rare overall, but were more frequent in the distal than in the proximal region in both CON21 (2.17% versus 1.16%) and KAP4 (2.51% versus 1.42%) (**Table S4**). Thus, although the proximal region showed the clearest enrichment of TATA-like sequence features, neither initiation region could be explained simply by a canonical TATA–Inr promoter configuration.

In contrast to the A/T-rich proximal region, the distal initiation region was enriched for a distinct set of motifs, many containing CACG/CACGTG-like sequence cores (**Figure S7**). These motifs were particularly prominent in KAP4, whereas CON21 showed fewer strongly enriched distal motifs and generally weaker enrichment signals. The major contrast therefore occurred not simply between the two lineages, but between the proximal and distal initiation regions themselves: the proximal region was preferentially associated with A/T-rich and TATA-like sequence features, whereas the distal region was characterized by CACG/CACGTG-like motifs and a modestly higher frequency of Inr-like sequences. Together, these results suggest that the double-peak transcription-initiation architecture is accompanied by spatially distinct core-promoter and *cis*-regulatory sequence environments. Rather than representing two equivalent initiation sites governed by the same promoter grammar, the proximal and distal peaks appear to be associated with different combinations of promoter elements, with additional lineage-associated quantitative differences in motif enrichment, particularly in the distal region.

### Promoter architecture and developmental expression dynamics

To further examine whether TSR architecture is associated with gene-expression differences, we compared expression levels of genes linked to broad, intermediate, and peaked promoters across developmental stages in KAP4 and CON21. In both lineages, a consistent and ordered pattern was observed: genes with broad promoters exhibited the highest expression levels, intermediate promoters showed intermediate expression, and peaked promoters were associated with the lowest expression levels across nearly all developmental stages (**Figure 5**). This pattern supports a general coupling between promoter shape and gene-expression levels, suggesting that promoter architecture reflects quantitative differences in transcriptional output.

**Figure 5.**
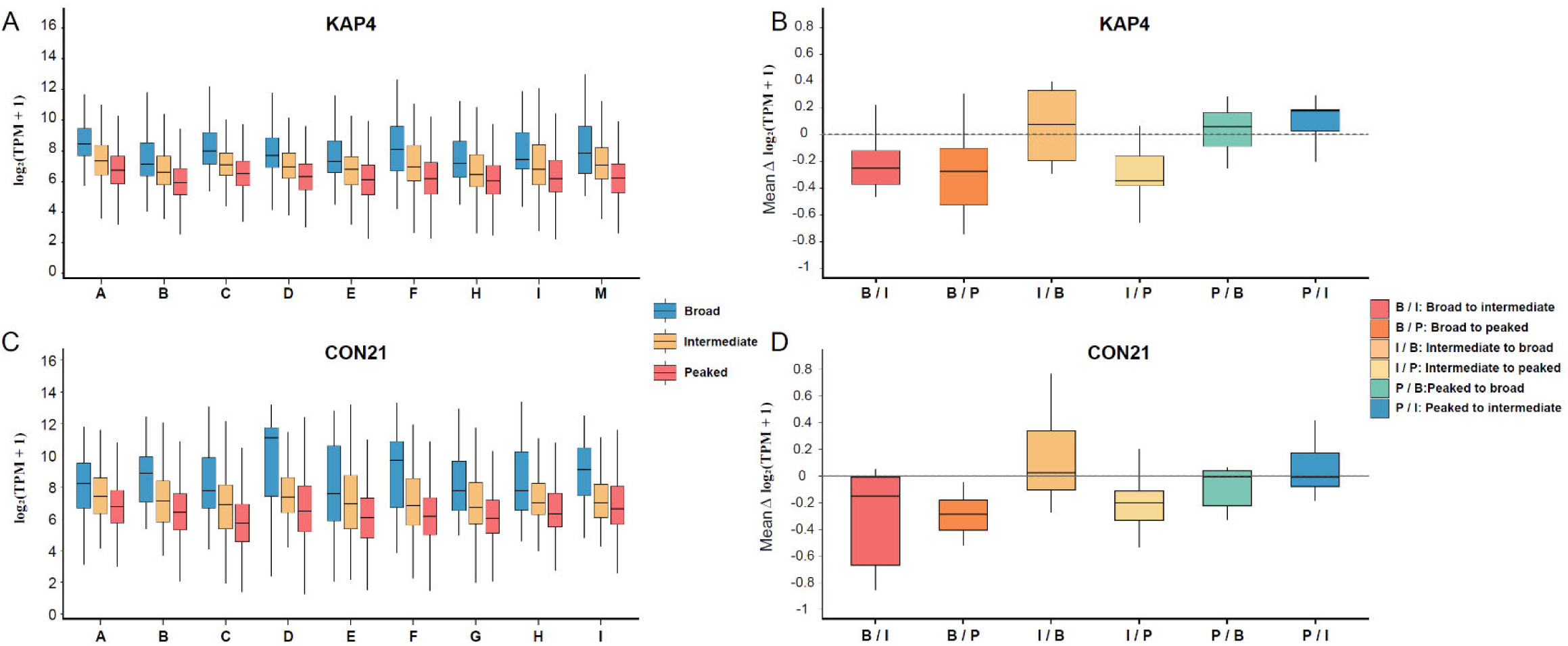
Association between promoter architecture and gene expression during development in KAP4 and CON21. Expression levels of genes associated with broad, intermediate, and peaked promoters across developmental stages in KAP4 (A) and CON21 (C). Gene expression is shown as log_2_(TPM + 1). (B, D) Changes in gene expression associated with promoter-type transitions between consecutive developmental stages in KAP4 (B) and CON21 (D). Expression change was calculated as the mean Δlog_2_(TPM + 1) between adjacent stages for genes that changed promoter type. The dashed horizontal line indicates no expression change. Negative values indicate decreased expression after promoter-type transition, whereas positive values indicate increased expression.

We next examined whether promoter-type switching within the same gene was associated with expression changes between consecutive developmental stages (**Figure S8**). In KAP4, transitions toward more peaked promoter states were consistently associated with reduced expression, including broad-to-intermediate, broad-to-peaked, and intermediate-to-peaked transitions, with mean decreases of approximately 13–17% (**Figure 5B**). In contrast, transitions toward broader promoter states were associated with only modest increases in expression, generally less than 10%. A similar but more pronounced pattern was observed in CON21, where transitions toward more peaked states resulted in expression decreases of approximately 17–34%, with the strongest effect observed for broad-to-peaked transitions (**Figure 5D**). Transitions toward broader states again produced only small increases, generally no more than ∼11%. Together, these results indicate an asymmetric relationship between promoter-shape switching and transcriptional output in both lineages, with transitions toward more peaked promoters associated with larger decreases in expression than the increases accompanying transitions toward broader promoters.

We then examined whether promoter-type divergence between KAP4 and CON21 was associated with expression divergence. For genes expressed in both lineages, we compared the absolute expression difference, measured as |Δlog_2_(TPM + 1)|, among genes with either conserved or divergent promoter types. Overall, genes with different promoter types between KAP4 and CON21 did not show consistently greater expression divergence than genes with the same promoter type (**Figure S9**). The distributions of expression-divergence largely overlapped among all promoter-pair categories. Promoter pairs representing different classes, such as Broad-Intermediate, Broad-Peaked, and Intermediate-Peaked, showed median values comparable to those of same-type promoter pairs, including Broad-Broad and Intermediate-Intermediate. Notably, genes classified as Peaked-Peaked tended to show relatively high expression divergence and a wider range of values, indicating that substantial expression differences can occur even when promoter shape is conserved between the two genetic backgrounds (**Figure S9**).

These results suggest that promoter-shape divergence alone, at least at the broad classification level used here, is not sufficient to explain the overall magnitude of expression divergence between KAP4 and CON21. This is also consistent with the motif analysis, where even promoters assigned to the same shape category in KAP4 and CON21 could exhibit divergent enriched motifs (**Figure 4C**). Thus, genes with the same broad promoter type may still differ substantially in their underlying regulatory architecture. Expression divergence is therefore likely influenced by additional regulatory features, including sequence-level variation within promoters, motif composition, transcription factor availability, chromatin context, and other *cis*-and *trans*-regulatory factors. However, when we focused specifically on genes showing greater than two-fold expression differences between KAP4 and CON21, promoter-type divergence became significantly associated with expression divergence. Genes with different promoter types were more likely to exhibit >2-fold expression differences than genes with conserved promoter types (χ² = 9.0962, *P* = 0.00256). This pattern suggests that although promoter-type divergence does not uniformly increase expression divergence across all shared genes, it may contribute to major expression shifts in a subset of genes. Together, these results indicate that promoter shape changes are not a simple one-to-one predictor of expression divergence, but they may represent one component of a broader regulatory mechanism underlying large expression differences between the two genetic backgrounds.

## Discussion

In this study, we generated an integrated developmental transcriptome and promoter atlas for two lineages within the *Daphnia pulex* species complex. By combining chromosome-level genome assemblies, RNA-seq, transcription start site profiling, and comparative evolutionary analyses, our study reveals how regulatory divergence can accumulate while the broader developmental program remains conserved. KAP4 and CON21 share most of their active protein-coding transcriptomes and broadly similar developmental trajectories, yet differ in the temporal deployment of genes and several features of transcription-initiation architecture. This combination is particularly informative because the two lineages are relatively closely related, with a mean synonymous divergence of only ∼4.7% among orthologous genes. Thus, developmental regulatory divergence in the *D. pulex* complex appears to involve quantitative remodeling of gene deployment and transcription initiation rather than wholesale restructuring of the underlying developmental program.

The two lineages exhibit broadly similar developmental trends in PCG expression, but differ quantitatively in the size and composition of their expressed transcriptomes. In both KAP4 and CON21, the number of expressed PCGs increased from early embryogenesis toward later developmental stages, consistent with the progressive activation and expansion of the zygotic developmental program. Despite this shared developmental trajectory, KAP4 consistently expressed more PCGs at most stages and showed a larger set of clone-specific expressed genes (**Figure 2A**). However, because a similar overall proportion of annotated PCGs was expressed in the two lineages, the greater absolute number of expressed genes in KAP4 should not necessarily be interpreted as evidence of a broader developmental program. Instead, this difference may partly reflect variation in gene annotation or lineage-specific gene expression.

The divergence between the two lineages was even more pronounced for lncRNAs. KAP4 expressed substantially more lncRNAs than CON21 across development, and a larger fraction of KAP4 lncRNAs showed persistent expression across stages **(Figure S1**). Rapid evolutionary turnover of lncRNA repertoires and transcription has been observed across animals, with many lncRNAs showing lineage-specific expression, whereas a smaller subset exhibits deeper evolutionary conservation and stronger sequence or expression constraint (Necsulea and Kaessmann 2014;; Necsulea et al. 2014; Hezroni et al. 2015). The greater lineage specificity observed here is therefore consistent with the broader view that noncoding transcription constitutes a comparatively dynamic component of regulatory evolution. However, this interpretation requires particular caution in *Daphnia*. lncRNAs are typically weakly expressed, their gene models are sensitive to transcript coverage and annotation procedures, and sequence divergence makes orthology substantially more difficult to establish than for protein-coding genes (Necsulea and Kaessmann 2014). Consequently, our data demonstrate extensive divergence in the observed lncRNA repertoires but cannot determine how much reflects true gain, loss, or regulatory turnover. Future work integrating Iso-Seq-supported transcript models, and chromatin accessibility, would help distinguish functional lncRNAs from incompletely annotated or weakly expressed transcripts.

Stage-specific genes provide a clearer connection between developmental expression and evolutionary constraint. These genes were expressed at substantially lower levels and exhibited elevated π_N_/π_S_ and d_N_/d_S_ relative to genes expressed throughout development. In *Drosophila*, narrower developmental expression breadth is associated with faster evolutionary rates, consistent with reduced pleiotropic constraint on temporally restricted genes (Artieri et al. 2009). The elevated evolutionary rates of stage-specific genes in *Daphnia* therefore likely reflect the combined effects of lower expression and reduced temporal pleiotropy. Importantly, their elevated π_N_/π_S_ and d_N_/d_S_ are more consistent with relaxed purifying selection than with widespread positive selection. Indeed, all stage-specific gene sets had neutrality index (NI) values greater than 1, indicating an excess of nonsynonymous polymorphism relative to divergence and further supporting reduced efficacy of purifying selection. Such reduced constraint may increase the evolutionary flexibility of stage-specific genes, allowing accumulation of nonsynonymous variation and potentially facilitating lineage-specific developmental functions, while also making these genes more prone to evolutionary turnover.

Comparative analyses of developmental expression profiles revealed that transcriptome divergence between KAP4 and CON21 is highly stage-dependent, and our data do not provide strong support for a canonical embryonic hourglass pattern. In *Drosophila* and several other systems, mid-development often shows enhanced transcriptomic conservation, although both the position and strength of this conserved period vary among taxa and analytical approaches (Kalinka et al. 2010; Roux et al. 2015; Drost et al. 2017). In our data, the strongest overall similarity between KAP4 and CON21 was observed during late embryonic and early juvenile stages, especially stages F and H, rather than during the middle of embryogenesis. We therefore interpret this pattern as a shifted or extended window of developmental conservation rather than as evidence against the hourglass principle itself.

The contrasting behavior of different similarity metrics further emphasizes that developmental conservation is multidimensional. Stage C illustrates this point particularly clearly. Although shared orthologs showed a high Pearson correlation in expression at this stage, indicating conservation of their relative expression patterns, the two lineages shared relatively fewer expressed genes and differed substantially in absolute expression levels. Thus, conservation of relative expression structure does not necessarily imply overall transcriptomic similarity. In contrast, stages F and H showed greater similarity across multiple measures, including expressed-gene overlap and expression magnitude. These results caution against defining developmental conservation from correlation alone and suggest that different components of the transcriptional program can diverge at different rates. Heterochronic differences between lineages may also contribute, because nominally equivalent morphological stages need not be perfectly aligned at the molecular level (Roux et al. 2015). The transition from late embryogenesis to independent juvenile life in *Daphnia* may additionally represent a period of strong functional integration, potentially imposing greater constraints on gene deployment.

The TSS data reveal a second level at which conserved organization coexists with evolutionary divergence. Both lineages showed a reproducible aggregate two-peak distribution of TSS signals upstream of annotated translation-start sites, but differed in the relative use of distal and proximal initiation regions. Because these peaks were obtained by aggregating signals across thousands of genes, they should not be interpreted as evidence that individual genes generally contain two TSSs at these positions. Instead, the distribution may reflect mixtures of genes with distinct dominant TSS positions, or promoter architectures. Alternative TSS usage is widespread in metazoans and represents an important layer of gene regulation (Alfonso-Gonzalez and Hilgers 2024). Comparative analyses in *Drosophila* further show that TSS position, promoter activity, and peak shape can diverge among closely related species, providing a potential source of regulatory evolution (Main et al. 2013). The KAP4–CON21 differences may therefore reflect evolutionary shifts in the relative deployment of alternative transcription-initiation modes.

The distinct motif environments associated with proximal and distal initiation regions further suggest that these modes of initiation are not equivalent. In metazoans, core promoters can differ substantially in their sequence architecture, and motifs such as TATA, Inr, MTE, and DPE have been associated with distinct promoter classes and regulatory programs, particularly in *Drosophila* and mammals (Rach et al. 2009; Hoskins et al. 2011; Zabidi et al. 2015; Haberle and Stark 2018). In *Daphnia*, A/T-rich and TATA-like features were enriched upstream of a subset of STRIPE-seq-defined TSSs, whereas a canonical Inr is only detected in ∼2% of the genes. These results suggest that some promoter features characterized in established metazoan model systems may also occur in *Daphnia*, but they also caution against assuming that promoter sequence organization is conserved uniformly across animal lineages. The CACG/CACGTG-like motifs enriched in distal regions are also notable, although motif enrichment alone is insufficient to assign specific transcription factors. More broadly, the contrasting motif compositions of proximal and distal regions support the idea that alternative initiation environments differ in their *cis*-regulatory organization.

Promoter shape was also closely associated with transcriptional output. In both lineages, genes with broad promoters showed higher average expression than genes with intermediate or peaked promoters (**Figure 5**). This pattern is consistent with observations in other metazoans, where broad or dispersed promoters are frequently associated with broadly expressed genes, whereas focused promoters are more often associated with regulated or cell-type-specific expression programs (Lenhard et al. 2012; Kedmi et al. 2021; Roeder 2023). Developmental and housekeeping regulation can also involve differences in core-promoter architecture and enhancer–promoter compatibility (Zabidi et al. 2015; Kedmi et al. 2021), suggesting that promoter architecture may influence how regulatory inputs are integrated. However, the relationship between promoter shape and expression level should be interpreted cautiously. Our expression measurements were obtained from whole animals and therefore represent averages across multiple tissues and cell types. Genes with relatively low overall expression may still be strongly expressed in restricted cell populations, a possibility that cannot be resolved with the present bulk data. Thus, the association between more peaked promoter states and lower average expression does not imply that these promoters are intrinsically weak, nor does it establish a simple causal relationship between promoter shape and transcriptional output. Promoter shape and promoter strength can also behave as partially independent regulatory properties (Schor et al. 2017). This distinction may help explain why promoter-type divergence was not uniformly associated with expression divergence between KAP4 and CON21 (**Figure 5B, 5D**). Broad promoter categories summarize only one dimension of regulatory architecture: promoters assigned to the same class may still differ in dominant TSS position, initiation strength, motif composition, enhancer interactions, or chromatin context. TSS positions and promoter usage can evolve among closely related species (Main et al. 2013), while promoter strength and shape can have partly separable regulatory architectures (Schor et al. 2017). The enrichment of promoter-type differences among genes with greater than twofold expression divergence therefore suggests that promoter remodeling is associated particularly with large expression shifts, while accounting for only part of genome-wide regulatory divergence.

Overall, our results reveal substantial remodeling of developmental transcription initiation between two *D. pulex* lineages despite their relatively modest sequence divergence. The central conclusion is not that promoter architecture alone explains developmental-expression divergence, but that transcription-initiation architecture can evolve detectably while the broader developmental transcriptional program remains conserved. Changes in TSS usage, promoter shape, and local *cis*-regulatory sequence composition therefore provide multiple routes for tuning transcription over relatively short evolutionary timescales. This combination of conserved developmental output and flexible transcription-initiation architecture may provide an important mechanism by which closely related lineages accumulate regulatory differences while maintaining developmental robustness.

## Materials and Methods

### *Daphnia* culture and developmental stage definition

The *Daphnia pulex* isolates KAP4 and CON21 were used as two clonal lineages in this study. KAP4 was originally collected from a temporary pond, Kickapond (40.122513, −87.735059), in the United States in spring 2013, whereas CON21 was collected from the Czech Republic (50.024578, 13.894197). Both isolates are cyclically parthenogenetic, but were maintained under laboratory conditions that favor parthenogenetic reproduction, allowing multiple genetically identical individuals from each clone to be collected for sequencing and downstream analyses.

Clonal cultures of KAP4 and CON21 were maintained in COMBO artificial medium (Kilham et al. 1998) at 18°C under a 16 h light:8 h dark photoperiod. Each culture was kept in a 150 mL glass beaker containing 80 mL of medium and was initiated with four adult females carrying parthenogenetic eggs in their brood chambers. This culture design maintained clonal reproduction while minimizing overcrowding and suppressing male production. Animals were fed daily with the green alga *Scenedesmus obliquus* at a final concentration of approximately 200,000 cells/mL. Cultures were transferred to fresh medium every two weeks to reduce waste accumulation. At each transfer, four gravid females carrying parthenogenetic eggs were randomly selected and placed into fresh beakers to maintain each clonal lineage.

Embryonic stages were identified based on morphological characteristics under a dissecting microscope. Embryonic development was subdivided into six stages: stage A (1–8 h), corresponding to eggs newly deposited in the brood chamber; stage B (9–18 h), marked by the appearance of a transparent edge; stage C (19–24 h), characterized by segmentation of the chorion; stage D (25–32 h), involving embryo elongation and the formation of a visible head; stage E (33–45 h), during which red eyes fill in the black pigment and an irregular heartbeat is present but regular movement is absent; and stage F (45–56 h), defined by the formation of a single black eye, regular heartbeat, and the capacity of embryos to swim freely when removed from the brood chamber. Juveniles were sampled at three time points after release from the brood chamber: 8 h (stage G, approximately instar 1), 56 h (stage H, approximately instar 2), and 104 h (stage I, approximately instar 4). Adults (stage M) were defined as individuals maintained for seven days after release from the brood chamber. Adult females were euthanized, and their eggs were carefully dissected from the brood chambers.

For embryonic stages, 70–120 eggs were collected for early stages (A–C), and 20–50 embryos were collected for later stages (D–F). To preserve RNA integrity and halt further development, 1 μL of RNAshield was added for every 15–25 eggs or embryos. Samples were kept on ice throughout dissection and subsequently stored at −80°C until RNA extraction. Juvenile samples were obtained from mothers carrying late stage-F embryos. These mothers were isolated from bulk cultures and transferred to fresh beakers until parturition. At each juvenile stage, 20–50 individuals were collected, preserved in RNAshield, and immediately frozen at −80°C. For adult samples, 20 females were collected seven days after release from the brood chamber. To minimize embryonic contamination, eggs or embryos were carefully removed from the brood chambers before adult tissue was preserved for RNA extraction.

### Genome Sequencing, Assembly, and Annotation

To reduce potential contamination from gut-associated bacteria and algae, individuals used for genome sequencing were maintained in fresh COMBO medium without food for 48 h before DNA extraction. For each clone, high-molecular-weight genomic DNA was extracted from approximately 250 clonal individuals of KAP4 and CON21 using the MasterPure™ Complete DNA and RNA Purification Kit (Lucigen, Cat. No. MC85200), following the manufacturer’s protocol. DNA samples were submitted to the University of California, Irvine, for PacBio HiFi library construction and sequencing. Additional live individuals were sent to Phase Genomics for Hi-C library construction and sequencing.

To provide transcript evidence for genome annotation, individuals from each clone were exposed to six abiotic perturbations associated with anthropogenic environmental stress: elevated temperature, low pH, UV light, nickel, atrazine, and sodium chloride. Except for UV exposure, all treatments were conducted for 24 h without additional algal food. Atrazine and nickel treatments were performed at final concentrations of 4 mg/L and 0.03 g/L, respectively. Additional treatments included exposure to 26°C, pH 5.0 medium, or medium supplemented with 5 g/L NaCl. UV treatments were conducted in 250 mL beakers containing 50 mL of medium, with beakers placed 10.5 cm below 30 W, 36-inch Reptisun 5.0 UVB fluorescent bulbs for 4 h. All treatments, except the 26°C temperature treatment, were conducted at 18°C. RNA was collected immediately after treatment. Total RNA was extracted using either the RNeasy Mini Kit or the Direct-zol™ RNA Miniprep Kit (Zymo Research, Cat. No. R2052). RNA samples were submitted to the Translational Genomics Research Institute in Phoenix for Illumina RNA-seq and to the University of California, Irvine, for Iso-Seq library construction and sequencing.

PacBio HiFi reads were first assembled using hifiasm (Cheng et al. 2021) with default parameters. The primary assemblies were then filtered to remove putative contaminants and assembly artifacts. Contigs with GC content greater than 50% were marked as potential bacterial or algal contaminants. In addition, each contig was searched against the NCBI non-redundant (NR) database, and sequences with best matches to bacterial or algal origins were excluded. To reduce the false retention of alternative haplotypes from heterozygous regions, redundant haplotypic contigs were further purged using purge_dups (Guan et al. 2020).

The filtered assemblies were scaffolded using Hi-C interaction data. Hi-C contact maps were generated using Juicer tools (Durand et al. 2016a), and scaffolds were visualized and manually curated using Juicebox v1.11.08 (Durand et al. 2016b). Repetitive sequences in the final genome assemblies were identified and masked using RepeatMasker v4.1.2-p1 (Smit et al. 2004) with default parameters. Genome completeness was evaluated using BUSCO with the Arthropoda odb12 dataset (Simão et al. 2015). Both genome assemblies were annotated using the NCBI Eukaryotic Genome Annotation Pipeline, EGAPx, incorporating transcriptomic evidence from RNA-seq and Iso-Seq data.

### Total RNA Isolation, Library Construction, and Sequencing

Total RNA was extracted from developmental samples using the Zymo Direct-zol RNA kit in combination with TRIzol reagent. For each sample, 300 μL of TRIzol reagent was added immediately before homogenization, and samples were homogenized for 5 min using a motor-driven grinder. RNA was then purified following the standard spin-column-based Direct-zol protocol, including a 15 min DNase I digestion step to remove residual genomic DNA. Immediately after extraction, RNA concentration was measured using a Thermo Fisher Qubit fluorometer. Samples with sufficient RNA yield were further assessed for RNA integrity using an Agilent TapeStation 4200. Purified RNA samples were stored at −80°C until library construction.

For transcription start site profiling, STRIPE-seq libraries were prepared following the protocol described by Policastro et al. (2020). To standardize input among samples, 100-150 ng of total RNA was used for each library. Terminator exonuclease (TEX) digestion was first performed to remove uncapped RNA molecules, followed by reverse transcription to capture the 5′ ends of capped mRNAs. Before PCR amplification, bead-based size selection was performed using a 0.8:1 ratio of RNA Clean XP beads to product to enrich for appropriately sized cDNA fragments. After PCR amplification, two additional size-selection steps, using bead-to-product ratios of 0.6:1 and 0.7:1, were performed to narrow the library fragment-size range to approximately 250–1,000 bp. STRIPE-seq library quality was evaluated using an Agilent TapeStation 4200. Libraries with sufficient concentration, defined as greater than approximately 750 pg/μL within the 250–2,000 bp size range, and with minimal signal below 250 bp, were selected for sequencing. Libraries showing substantial low-molecular-weight products below 250 bp were subjected to additional bead-based size selection before sequencing to remove incomplete or short fragments.

For RNA-seq, the remaining RNA from the same developmental samples was used for standard RNA-seq library preparation after STRIPE-seq library construction. This design allowed gene-expression profiling and transcription start site profiling to be generated from matched developmental samples, enabling integration of RNA-seq-based expression estimates with STRIPE-seq-based promoter analyses. Both embryonic STRIPE-seq and RNA-seq libraries were sequenced at TGen using an Illumina NovaSeq 6000 platform.

### RNA-seq Read Processing, Expression Quantification, and Stage-Specific Expression Analysis

Raw RNA-seq reads from each biological replicate were processed independently throughout quality filtering, read mapping, and expression quantification. Reads were first filtered using Trimmomatic v0.36 (Bolger et al. 2014) with the parameters AVGQUAL:20, ILLUMINACLIP:TruSeq3-PE.fa:2:30:10, TRAILING:20, and MINLEN:50. To remove reads derived from ribosomal RNA (rRNA), the trimmed reads were aligned against a *Daphnia pulex* rRNA reference, and read pairs that did not map to the rRNA sequences were retained using SAMtools (Li et al. 2009). The resulting non-rRNA reads from KAP4 and CON21 were separately aligned to their corresponding genome assemblies using HISAT2 (Kim et al. 2019). Alignment files were subsequently processed using SAMtools, and only uniquely mapped reads with a mapping quality score of 60 were retained for expression quantification.

Gene-expression levels were quantified independently for each biological replicate using StringTie (Pertea et al. 2015, 2016), with the corresponding genome annotation for each lineage. Expression levels were reported as transcripts per million (TPM), which accounts for both sequencing depth and transcript length and was used here to describe relative expression levels and developmental expression patterns (Wagner et al. 2012). TPM was calculated by first normalizing read counts by transcript length in kilobases to obtain reads per kilobase (RPK), followed by normalization of each gene’s RPK value to the sum of RPK values across all genes in the sample and multiplication by 10^6^. Biological replicates were not pooled before read mapping or expression quantification.

To distinguish reproducible expression from low-level transcriptional noise, expression was first evaluated at the level of individual biological replicates. Protein-coding genes (PCGs) were considered expressed at a developmental stage when their mean TPM across the three biological replicates was >1 and at least two of the three replicates individually had TPM >1. Because lncRNAs are generally expressed at substantially lower levels than PCGs, a threshold of TPM >0.2 was used for lncRNAs, with the same requirement that at least two of the three biological replicates exceeded this threshold. Stage-specific PCGs were defined as genes satisfying the expression criterion described above in only one developmental stage, while remaining below the TPM = 1 threshold in all other stages. Specifically, the mean TPM was required to be <1 in every non-focal stage, with at least two of the three biological replicates also having TPM <1. Stage-specific lncRNAs were identified using the same procedure with a TPM threshold of 0.2.

Reproducibility among biological replicates was further evaluated using the coefficient of variation (CV), calculated as the standard deviation divided by the mean expression level across the three replicates within each developmental stage. Genes showing highly variable expression among biological replicates (CV >0.8) were excluded from subsequent analyses. This filtering removed 1.9% and 2.2% of genes from KAP4 and CON21, respectively. After replicate-level quality control and filtering, the mean TPM across the three biological replicates was used as the representative expression value for each gene at each developmental stage in downstream analyses.

### STRIPE-seq Read Processing and Identification of Transcription Start Sites

Raw STRIPE-seq reads from each biological replicate were processed independently using a pipeline adapted from the original STRIPE-seq protocol (Policastro et al. 2020). Reads were quality filtered and trimmed to remove Illumina adapters and low-quality bases using Trimmomatic v0.33 (Bolger et al. 2014). Reads containing the STRIPE-seq-specific adapter sequence (TATAGGG) and the immediately preceding 8-nt unique molecular identifier (UMI) were identified, after which the adapter and UMI sequences were removed while retaining UMI information for subsequent duplicate removal. Residual ribosomal RNA (rRNA)-derived reads were removed by alignment against a *Daphnia* rRNA reference using HISAT2 v2.1.0 (Kim et al. 2019). The remaining reads from each lineage were then aligned to the corresponding *Daphnia pulex* genome assembly using HISAT2. Alignment files were processed using SAMtools v1.8 (Danecek et al. 2021), and only uniquely mapped reads with a mapping quality score of 60 were retained for downstream analyses.

PCR duplicates were removed independently for each biological replicate. Reads sharing both the same genomic 5′-end position and identical UMI sequence were considered PCR duplicates and collapsed into a single molecule. The genomic 5′-end positions of the resulting non-redundant reads were used to define candidate transcription start sites (CTSSs). CTSSs were subsequently associated with annotated genes according to their genomic positions and strand orientations, enabling downstream characterization of transcription initiation and promoter architecture.

Processed CTSS data from individual biological replicates were imported separately into CAGEr v2.14.0 (Haberle et al. 2015). Tag counts were normalized independently for each replicate using the power-law normalization implemented in normalizeTagCount. To reduce the contribution of low-abundance transcription-initiation signals, only CTSSs with normalized abundance >2 TPM and complete chromosome, coordinate, and strand information were retained. Within each biological replicate, retained CTSSs located on the same strand and separated by ≤20 bp were clustered into transcription start regions (TSRs) using the distance-based clustering algorithm implemented in CAGEr (distclu, maxDist = 20). CTSSs on opposite strands were clustered independently, and singleton clusters were excluded (keepSingletonsAbove = 0). TSRs were therefore identified independently in each biological replicate before replicate-level integration.

To ensure that downstream promoter analyses were based on reproducible transcription-initiation signals, replicate-specific TSRs from the same developmental stage were compared according to their genomic positions and strand orientation. A TSR was considered reproducible and retained for downstream analyses only when it was detected in at least two biological replicates within the same developmental stage. Overlapping replicate-specific TSRs representing the same local transcription-initiation region were merged to generate a consensus TSR for that developmental stage. The resulting consensus TSR coordinates were subsequently applied to individual biological replicates so that CTSS distributions and promoter shape could be quantified over the same genomic interval across replicates.

Because Shape Index (SI) distributions differed among developmental stages, promoter shapes were classified using Gaussian mixture models (GMMs) implemented in the R package mclust v6.1.2 (Scrucca et al. 2016), rather than using fixed SI thresholds. Two- and three-component models were fitted to the SI distributions and compared using the Bayesian Information Criterion (BIC). The three-component model consistently showed higher BIC values and was therefore selected for downstream analyses. The three Gaussian components were ordered according to their mean SI values and designated Broad, Intermediate, and Peaked promoters from lowest to highest mean SI. Each TSR was assigned to the component with the highest posterior probability. Only classifications with a maximum posterior probability ≥0.95 were retained; TSRs below this threshold were classified as unclassified.

To assess the reliability of promoter-shape classification, five high-confidence Broad and five high-confidence Peaked TSRs were randomly selected from each sample, and their CTSS tag distributions were visually inspected. Broad promoters showed dispersed initiation signals across multiple genomic positions, whereas Peaked promoters showed concentrated signals at one or a few neighboring positions, consistent with the GMM-based classifications.

To assess whether aggregate TSS-position distributions were disproportionately influenced by genes with high transcription-initiation activity, we performed a gene-level normalization analysis. For each gene, CTSS counts within the analyzed upstream region were first summed to obtain the total CTSS abundance for that gene. The count at each CTSS position was then divided by this gene-level total, such that the normalized CTSS weights for each gene summed to 1. This normalization ensured that each gene contributed equally to the aggregate positional profile regardless of its absolute transcription-initiation activity. Normalized CTSS weights were subsequently summed across genes within each developmental stage for each positional bin, and the resulting values were divided by the total summed weight across all bins to obtain the relative positional frequency. As an additional sensitivity analysis, positional distributions were recalculated after excluding genes in the upper 5% of total CTSS abundance within each developmental stage. Only CTSSs located within 300 bp upstream of the annotated translation start site were included in this analysis.

### Calculation of Stage-wise Transcriptomic Divergence Metrics

To quantify developmental transcriptome divergence between KAP4 and CON21, we compared gene-expression profiles at corresponding developmental stages using one-to-one orthologous protein-coding genes identified between the two lineages. For each developmental stage, expression values were averaged across biological replicates before comparison. Gene expression was represented as log_2_(TPM + 1), where the log transformation reduces the influence of highly expressed genes and the addition of 1 allows genes with zero TPM values to be included.

We first compared the overlap of expressed genes between KAP4 and CON21 at each developmental stage. Protein-coding genes with TPM > 1 were considered expressed. The number of shared expressed genes was calculated as the intersection of the expressed orthologous gene sets in the two lineages. To account for differences in the total number of expressed genes among stages and lineages, we also calculated the Jaccard index:

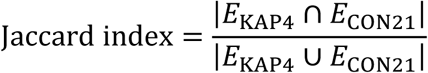

where E_KAP4 and E_CON21 represent the sets of expressed orthologous genes in KAP4 and CON21, respectively. A Jaccard index closer to 1 indicates greater overlap in the expressed gene repertoire, whereas a value closer to 0 indicates lower overlap.

We next quantified similarity in expression levels across orthologous genes using correlation-based metrics. Pearson correlation coefficients were calculated to measure the linear correspondence in log_2_(TPM + 1)-transformed expression values between KAP4 and CON21. Spearman rank correlation coefficients were also calculated to assess whether the relative ranking of gene-expression levels was conserved between the two lineages, without assuming a linear relationship. In addition, we calculated the proportion of orthologous genes showing expression differences within twofold between KAP4 and CON21. Genes were considered to be within twofold difference when the absolute difference in log_2_(TPM + 1) values between the two lineages was ≤ 1.

To quantify absolute expression divergence, we calculated mean absolute error (MAE), root mean squared error (RMSE), and median absolute log_2_ expression difference using log_2_(TPM + 1)-transformed expression values. MAE measures the average absolute expression difference between the two lineages and was calculated as:

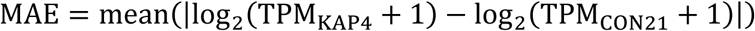

RMSE gives greater weight to genes with large expression differences and was calculated as:

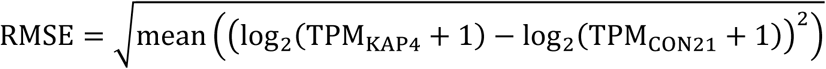

The median absolute log₂ expression difference was calculated as the median of

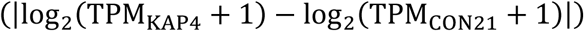

across orthologous genes at each stage.

### Motif Identification

Motif-enrichment analyses were performed to identify *cis*-regulatory sequence features associated with Broad, Intermediate, and Peaked promoter architectures. Promoter classes were defined independently for each sample using the Gaussian mixture model (GMM)-based classification described above.

For each TSR, the CTSS with the highest transcription-initiation signal within the TSR, was obtained from the CAGEr output. To ensure direct comparability among promoter classes and avoid biases associated with variable sequence lengths, a fixed promoter window extending from 100 bp upstream to 50 bp downstream of the dominant TSS (−100 to +50 bp) was used for motif analysis. Genomic intervals were generated in a strand-aware manner, and promoter sequences were extracted from the corresponding reference genome using bedtools getfasta (Quinlan 2014) with the strand-specific option (-s), such that sequences associated with negative-strand promoters were reverse complemented.

Within each sample, duplicated genomic intervals were removed before motif analysis so that each promoter sequence was represented only once. Promoters classified as Unclassified were excluded from the primary motif-enrichment analysis because of the uncertainty in their promoter-shape assignments, but were retained for sensitivity analyses where appropriate.

Transcription-factor binding motifs were obtained from the non-redundant JASPAR 2024 CORE insect collection (Rauluseviciute et al. 2024) and used in MEME format for downstream analyses. Motif enrichment was assessed using AME (Analysis of Motif Enrichment) implemented in MEME Suite v5.5.4 (McLeay and Bailey 2010; Bailey et al. 2015). Analyses were performed independently for each sample using a one-versus-rest design. For each focal promoter class, sequences assigned to that class were used as the foreground set, whereas sequences from the other two high-confidence promoter classes were combined as the control set. Thus, Broad, Intermediate, and Peaked promoters were each tested independently against the remaining promoter classes. Motif enrichment was evaluated using the one-tailed Wilcoxon rank-sum test based on sequence-level motif scores. Multiple-testing correction was applied across all tested motifs, and motifs with an adjusted *P* value < 0.05 were considered significantly enriched.

To reduce stage-specific sampling effects and avoid pseudoreplication of promoters repeatedly detected across development, motif enrichment was evaluated separately for each developmental sample rather than after pooling promoter sequences across stages. Motifs showing consistent enrichment in the same promoter class across multiple developmental stages were considered robust promoter-class-associated candidates. Enrichment patterns were subsequently compared between KAP4 and CON21 to identify conserved and lineage-associated differences in promoter *cis*-regulatory architecture.

### GO enrichment analysis for stage-specific expression

Functional descriptions were obtained by querying these genes against the NCBI non-redundant (nr) database, and Gene Ontology (GO) terms were assigned using Blast2GO (Conesa et al. 2005). KEGG pathway identifiers (e.g., K01312) were retrieved via the KAAS server (Moriya et al. 2007) (https://www.genome.jp/tools/kaas/) using the GhostX search program for fast and accurate protein homology searches.

To identify functional roles of genes with stage-specific expression patterns, we conducted Gene Ontology (GO) enrichment analysis. The enrichment analysis was performed using a χ² test implemented via the Perl module Statistics::ChisqIndep to assess overrepresented GO terms among these genes. To reduce redundancy and overlap among enriched terms, we used the web tool REVIGO (Supek et al. 2011) (http://revigo.irb.hr/) with the allowed similarity set to 0.8, while keeping all other parameters at their default values. This clustering approach grouped related GO terms and retained only the most representative ones. Statistical significance of enrichment was evaluated using the Benjamini-Hochberg method to correct for multiple testing, controlling the false discovery rate (FDR) at 5%.

### Data accession

The *Daphnia pulex* (KAP4) genome assembly is available at NCBI under accession GCF_021134715.1, and the CON21 genome assembly and its corresponding gene annotation are available from the European Nucleotide Archive (ENA) under accession GCA_988224025.1. The genome assembly for *D. obtusa* is available at GenBank under accession number JBOZXD000000000.1. All RNA-seq and STRIPE-seq data generated across developmental stages have been submitted to NCBI under study accession number PRJNA1332392.

## Acknowledgments

This work was supported by the National Natural Science Foundation of China (Grant No. 32471695) awarded to Z.Y.; the NIH (Grant R35-GM122566-01 to M.L.); the NSF Enabling Discovery through GEnomics (EDGE) program (Grant IOS-1922914 to M.L. and Andrew Zelhof, Indiana University).

