## Supplementary Tables and Figures for "Promoter architecture and developmental-expression divergence in the *Daphnia pulex* complex"

**Table S1.** Comparison of genome assembly and annotation statistics between *D. pulex* CON21 and KAP4 genome assemblies. PCGs, protein-coding genes; lncRNAs, long non-coding RNAs.

|  | KAP4 | | | | CON21 | | | |
| --- | --- | --- | --- | --- | --- | --- | --- | --- |
| Chr. | Size (Mb) | # Contig | PCGs | lncRNA | Size (Mb) | # Contig | PCGs | lncRNA |
| Chr.1 | 8.3 | 1 | 1,029 | 200 | 8.6 | 1 | 998 | 150 |
| Chr.2 | 13.4 | 2 | 1,325 | 363 | 14.2 | 1 | 1,267 | 223 |
| Chr.3 | 13.3 | 1 | 1,386 | 355 | 12.4 | 1 | 1,242 | 251 |
| Chr.4 | 9.2 | 2 | 969 | 255 | 9.7 | 1 | 947 | 116 |
| Chr.5 | 12.0 | 1 | 1,257 | 408 | 13.3 | 1 | 1,208 | 262 |
| Chr.6 | 7.3 | 1 | 863 | 226 | 7.8 | 1 | 901 | 143 |
| Chr.7 | 12.3 | 1 | 1,657 | 446 | 13.8 | 1 | 1,634 | 228 |
| Chr.8 | 14.2 | 2 | 1,428 | 371 | 15.4 | 1 | 1,199 | 248 |
| Chr.9 | 9.6 | 1 | 1,162 | 410 | 11.8 | 1 | 1,152 | 239 |
| Chr.10 | 16.3 | 1 | 2,101 | 495 | 18.0 | 1 | 2,076 | 331 |
| Chr.11 | 6.6 | 1 | 945 | 270 | 8.0 | 1 | 964 | 218 |
| Chr.12 | 10.7 | 2 | 1,160 | 333 | 10.8 | 5 | 904 | 139 |
| Sum | 133.2 | 17 | 15,282 | 4,132 | 143.8 | 16 | 14,492 | 2,548 |

**Table S2.** Pearson correlation coefficients among biological replicates across different developmental stages were calculated using log₂^(TPM + 1)^-transformed expression values, where TPM represents transcripts per million.

|  | *KAP4* | | *CON21* | | | |
| --- | --- | --- | --- | --- | --- | --- |
| rep1 | rep2 | r | | rep1 | rep2 | r |
| A1 | A2 | 0.977 | | A1 | A2 | 0.992 |
| A1 | A3 | 0.933 | | A1 | A3 | 0.984 |
| A2 | A3 | 0.906 | | A2 | A3 | 0.991 |
| B1 | B2 | 0.961 | | B1 | B2 | 0.995 |
| B1 | B3 | 0.987 | | B1 | B3 | 0.989 |
| B2 | B3 | 0.956 | | B2 | B3 | 0.991 |
| C1 | C2 | 0.988 | | C1 | C2 | 0.924 |
| C1 | C3 | 0.986 | | C1 | C3 | 0.951 |
| C2 | C3 | 0.988 | | C2 | C3 | 0.982 |
| D1 | D2 | 0.992 | | D1 | D2 | 0.983 |
| D1 | D3 | 0.980 | | D1 | D3 | 0.986 |
| D2 | D3 | 0.988 | | D2 | D3 | 0.991 |
| E1 | E2 | 0.987 | | E1 | E2 | 0.971 |
| E1 | E3 | 0.973 | | E1 | E3 | 0.977 |
| E2 | E3 | 0.988 | | E2 | E3 | 0.989 |
| F1 | F2 | 0.973 | | F1 | F2 | 0.913 |
| F1 | F3 | 0.978 | | F1 | F3 | 0.945 |
| F2 | F3 | 0.970 | | F2 | F3 | 0.843 |
| G1 | G2 | 0.972 | | G1 | G2 | 0.983 |
| G1 | G3 | 0.990 | | G1 | G3 | 0.992 |
| G2 | G3 | 0.978 | | G2 | G3 | 0.980 |
| H1 | H2 | 0.950 | | H1 | H2 | 0.941 |
| H1 | H3 | 0.920 | | H1 | H3 | 0.941 |
| H2 | H3 | 0.927 | | H2 | H3 | 0.967 |
| I1 | I2 | 0.938 | | I1 | I2 | 0.935 |
| I1 | I3 | 0.949 | | I1 | I3 | 0.958 |
| I2 | I3 | 0.969 | | I2 | I3 | 0.956 |
| M1 | M2 | 0.946 | | M1 | M2 | 0.931 |
| M1 | M3 | 0.912 | | M1 | M3 | 0.909 |
| M2 | M3 | 0.901 | | M2 | M3 | 0.976 |
| Average | | 0.962 | | Average | | 0.962 |

**Table S3.** Summary of transcription start site (TSS) mapping in protein-coding genes of KAP4 and CON21 across developmental stages. The table lists the number of genes with identified TSSs≤1 kb, and their corresponding ratio to the total number of expressed genes.

| **Stage** | **Gene with TSS** | **Ratio to total expressed** | **Genes_with_2_TSRs** |
| --- | --- | --- | --- |
| KAP4 |  |  |  |
| A | 6,217 | 0.68 | 289 |
| B | 8,667 | 0.86 | 392 |
| C | 7,699 | 0.77 | 269 |
| D | 8,779 | 0.81 | 234 |
| E | 7,659 | 0.66 | 343 |
| F | 9,857 | 0.81 | 201 |
| H | 9,889 | 0.81 | 230 |
| I | 8,765 | 0.74 | 180 |
| M | 8,042 | 0.67 | 221 |
| **mean** | **8,397** | **0.76** | **262** |
| CON21 |  |  |  |
| A | 5,324 | 0.64 | 172 |
| B | 7,768 | 0.93 | 198 |
| C | 7,088 | 0.90 | 248 |
| D | 7,805 | 0.84 | 83 |
| E | 8,171 | 0.82 | 150 |
| F | 8,316 | 0.78 | 137 |
| G | 6,461 | 0.58 | 153 |
| H | 7,980 | 0.70 | 79 |
| I | 4,492 | 0.38 | 72 |
| **mean** | **7,045** | **0.73** | **144** |

**Table S4.** Distribution of Inr-like and TATA-like promoter motifs in proximal and distal regions of CON21 and KAP4. Sequence Count indicates the total number of promoter sequences analyzed in each region. Inr-like and TATA-like indicate the numbers of sequences containing the corresponding promoter motifs, and Ratio (%) represents the percentage of motif-containing sequences relative to the total number of sequences analyzed in each region.

| Lineage | Region | #Sequence | Inr-like | Ratio (%) | TATA-like | Ratio (%) |
| --- | --- | --- | --- | --- | --- | --- |
| CON21 | proximal | 2,328 | 27 | 1.16 | 251 | 10.78 |
|  | distal | 2,677 | 58 | 2.17 | 196 | 7.32 |
| KAP4 | proximal | 2,887 | 41 | 1.42 | 276 | 9.56 |
|  | distal | 5,857 | 147 | 2.51 | 369 | 6.30 |

**Figure S1.** Developmental gene expression intersections in KAP4 and CON21. UpSet plots showing the number of unique and shared transcribed protein-coding and lncRNA genes across ten developmental stages of *Daphnia pulex*. Only the twenty most frequent gene groupings are shown. Horizontal bars show the total number of genes expressed in each stage, while vertical bars indicate the number of genes shared across the connected stages (black dots).

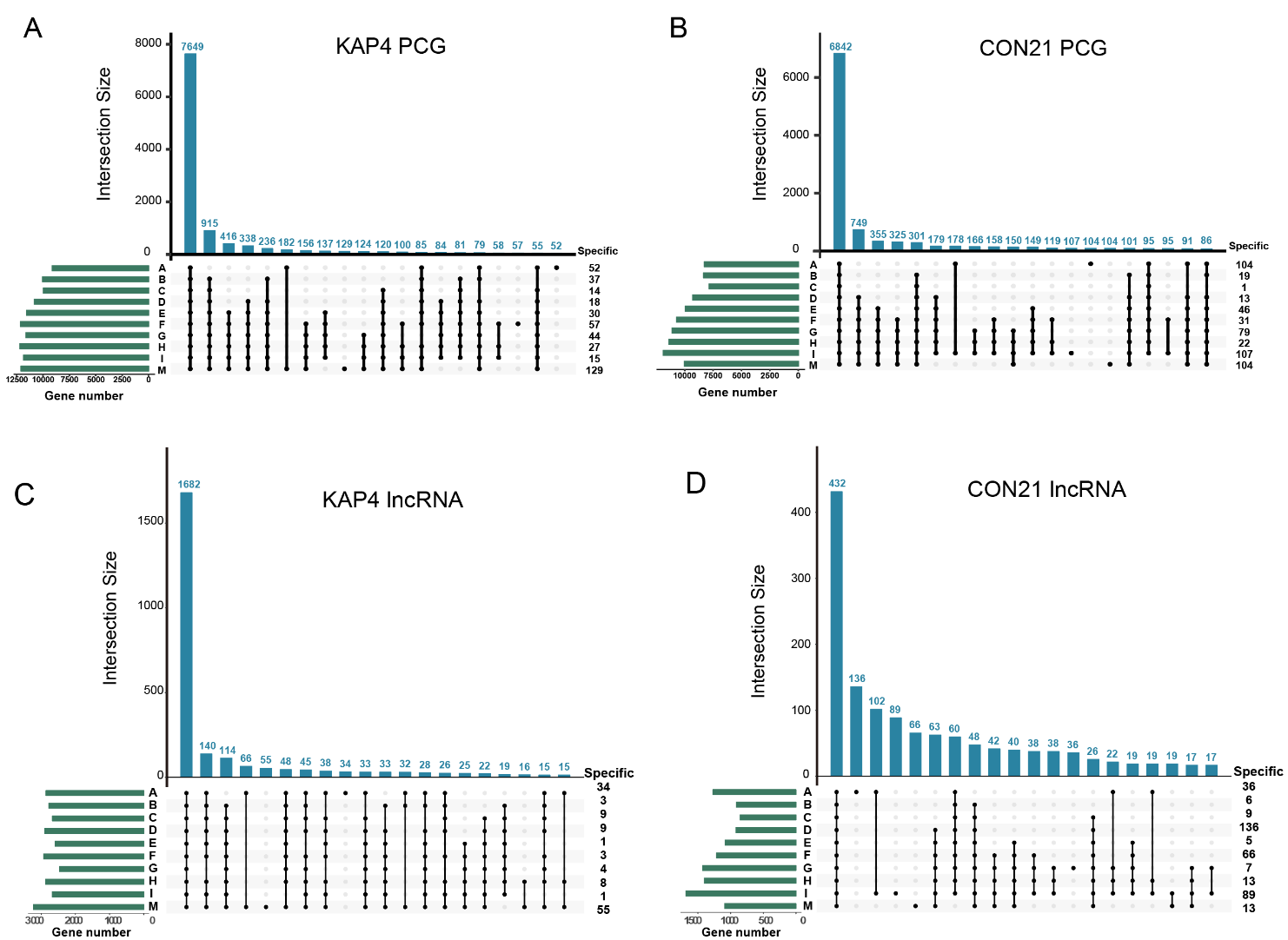

**Figure S2.** Venn diagram showing the numbers of expressed lncRNAs shared between KAP4 and CON21 or expressed only in one lineage at each developmental stage.

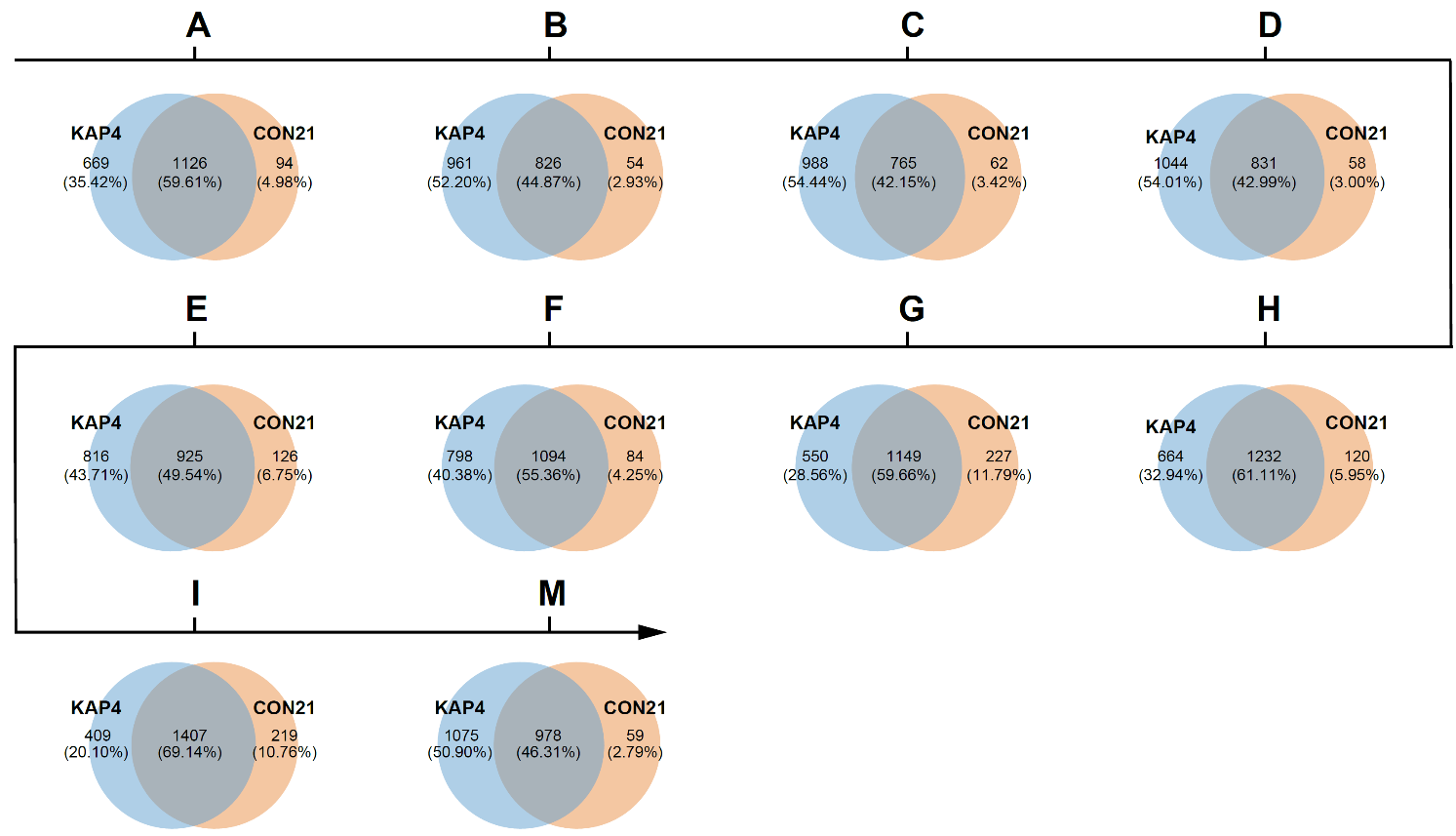

**Figure S3.** Functional enrichment analyses for KAP4 protein-coding genes expressed across all developmental stages. The top 30 enriched GO categories are shown.

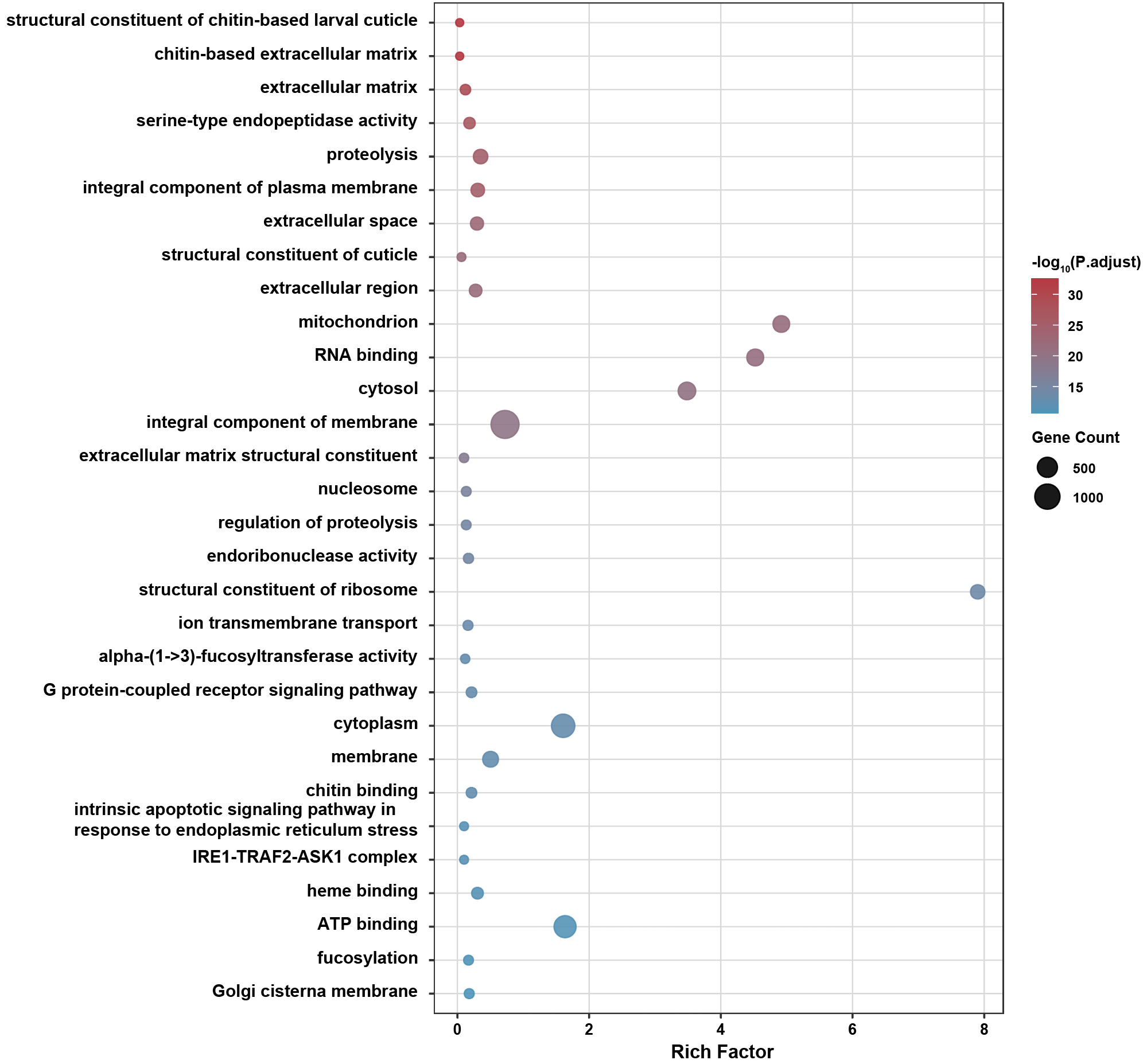

**Figure S4.** Functional enrichment analyses for CON21 protein-coding genes expressed across all developmental stages. The top 30 enriched GO categories are shown.

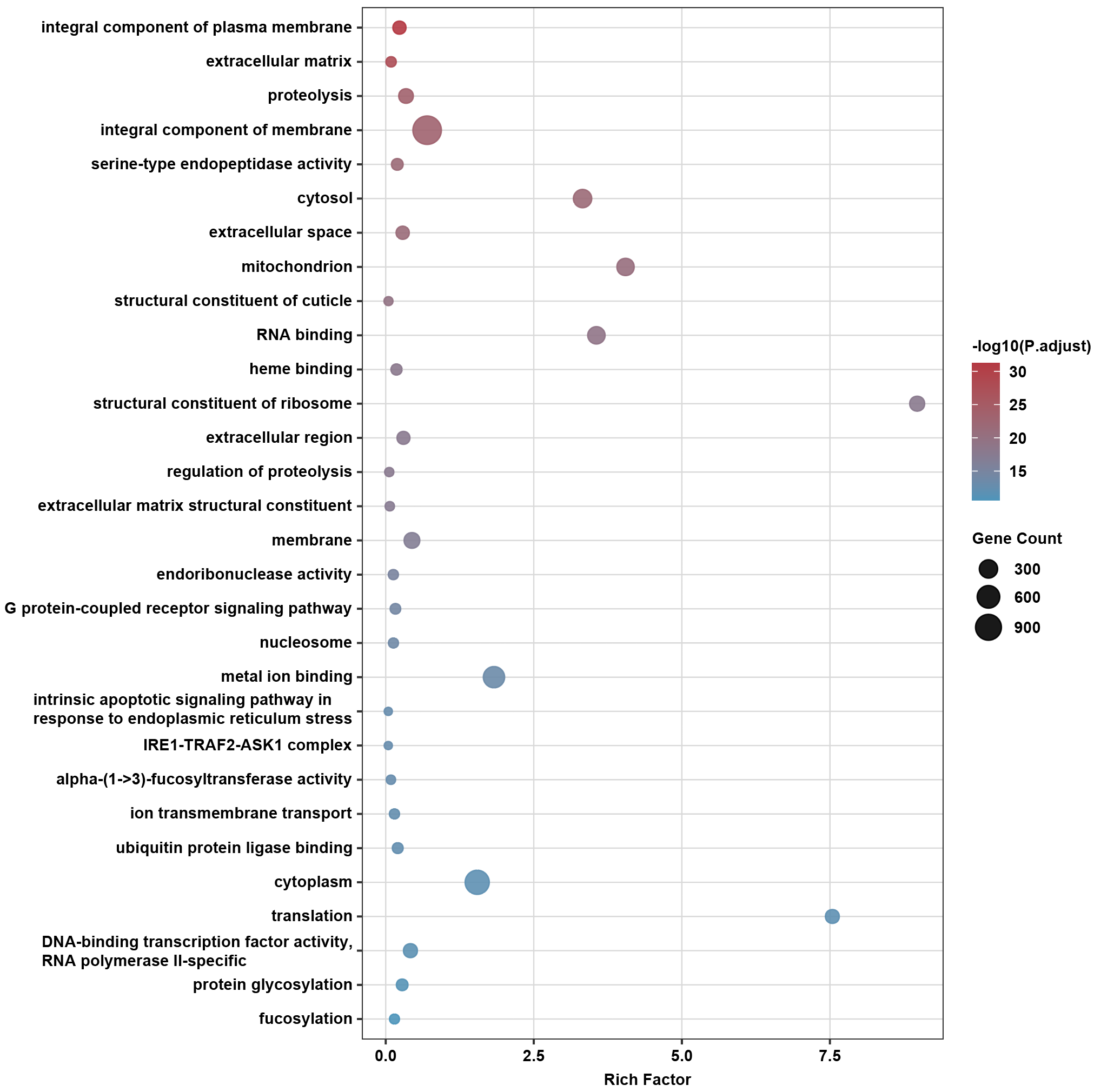

**Figure S5.** Functional enrichment analysis of KAP4 protein-coding genes with stage-specific expression. Enriched GO categories are shown separately for each developmental stage.

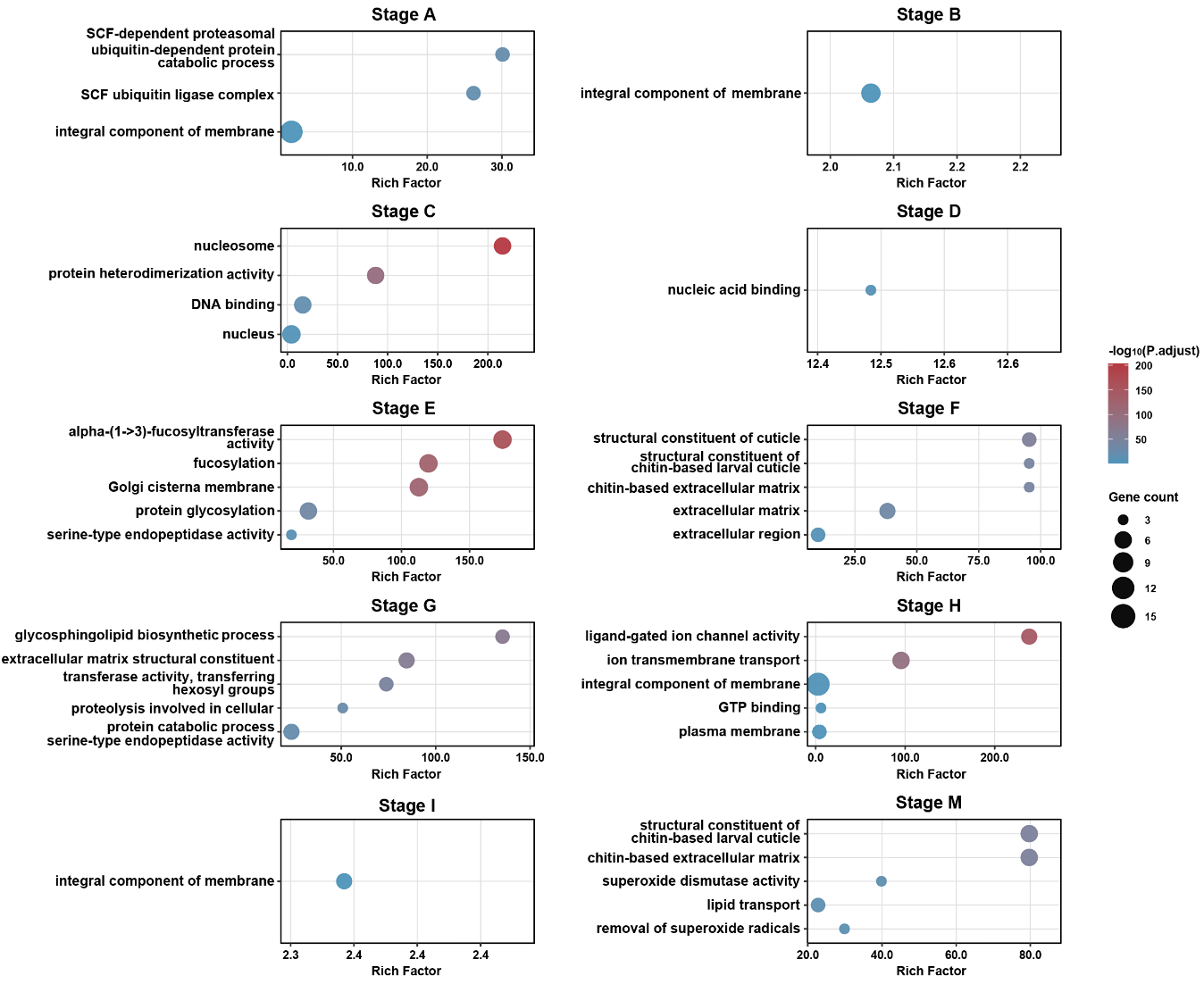

**Figure S6.** Functional enrichment analysis of CON21 protein-coding genes with stage-specific expression. Enriched GO categories are shown separately for each developmental stage.

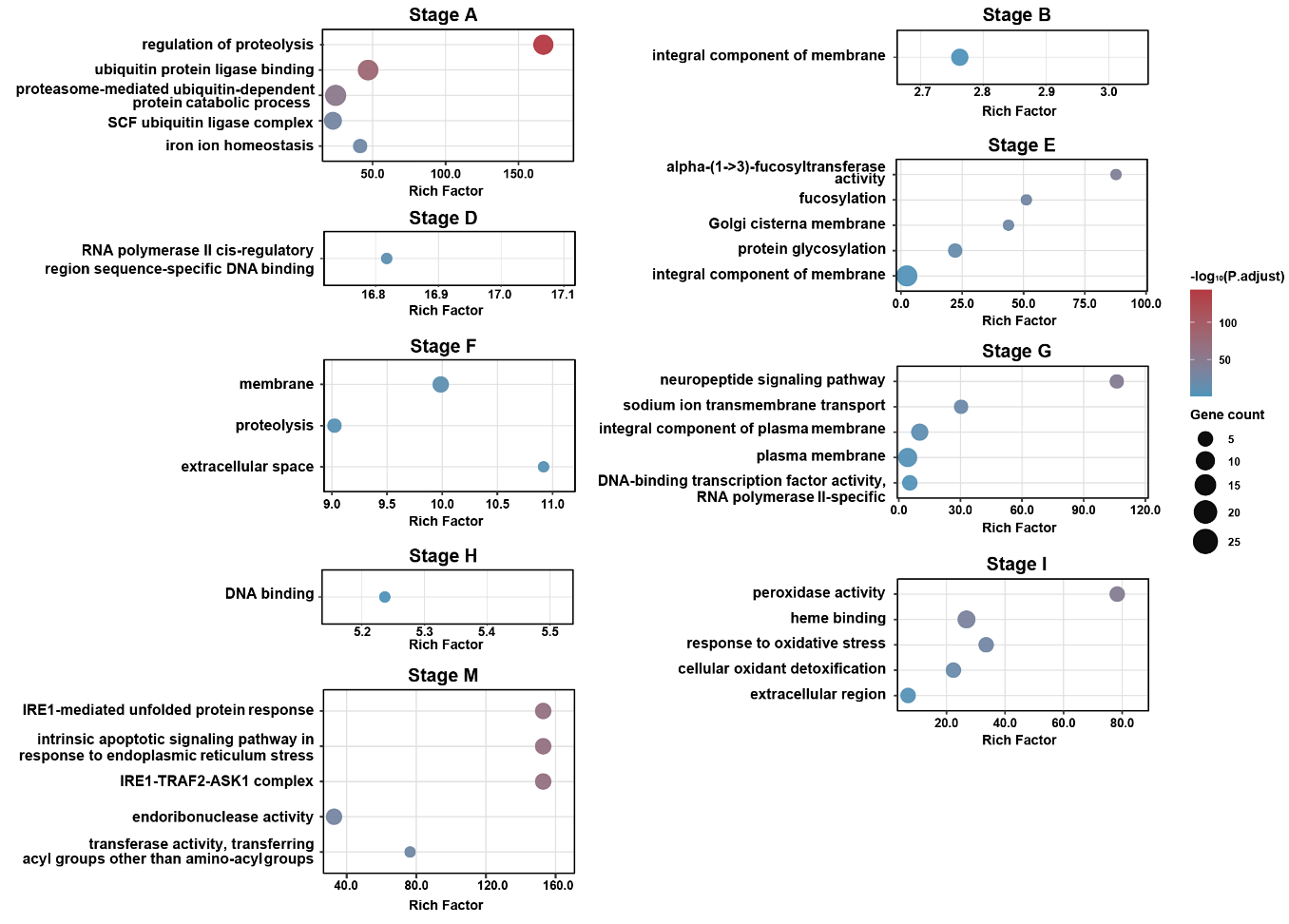

**Figure S7.** Motif enrichment in proximal and distal transcription initiation regions. Motif enrichment was analyzed separately in the proximal initiation region (-40 to -10 bp relative to the annotated translation start site) and the distal initiation region (-120 to -70 bp) in KAP4 and CON21. Only the most significantly enriched motifs (top 10 in each category based on −log_10_^FDR^) are shown to highlight major regulatory signals. Dot size indicates the proportion of target sequences containing each motif, and color intensity represents enrichment significance, shown as −log_10_^FDR^.

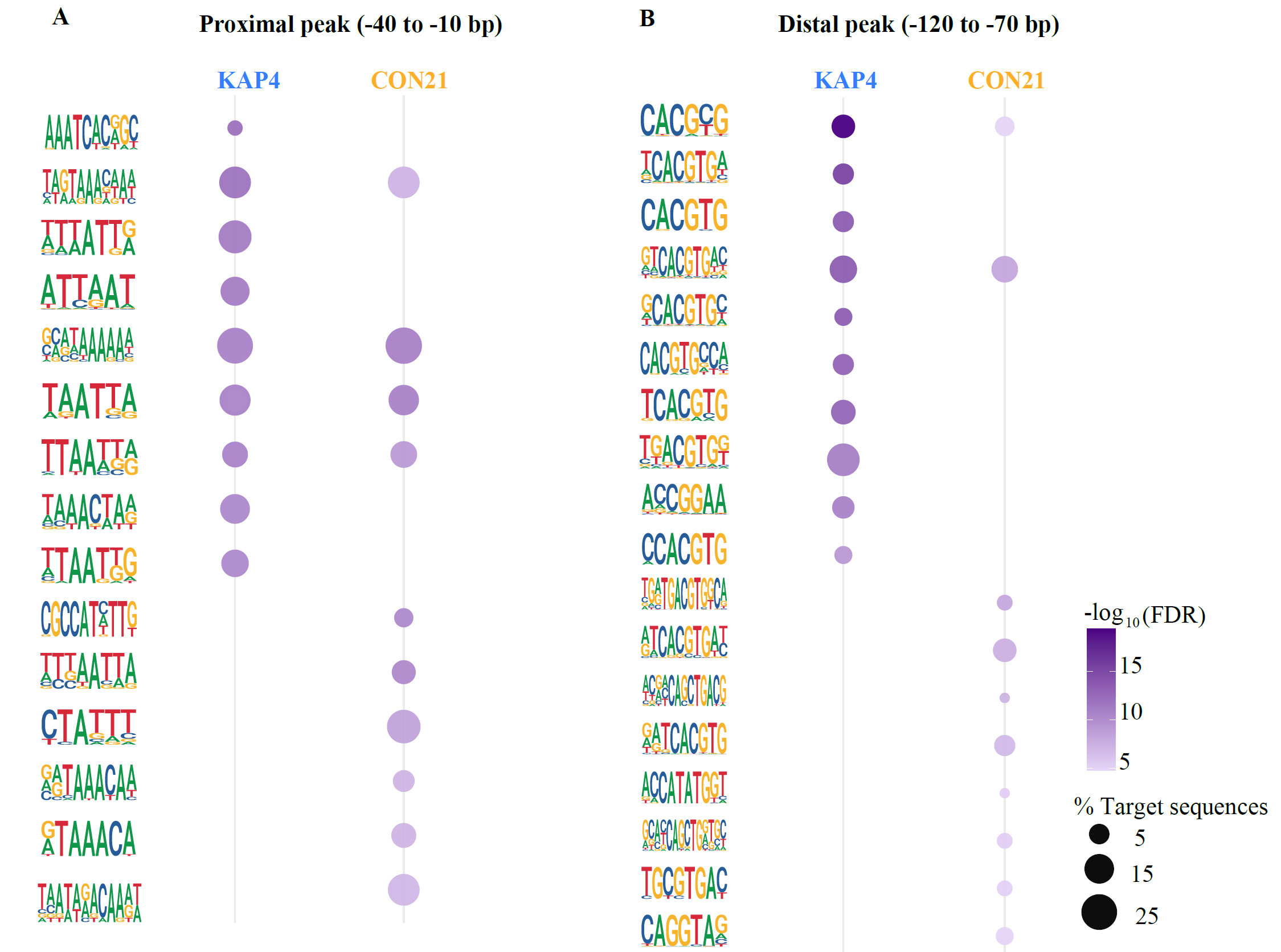

**Figure S8**. Developmental promoter-type transitions in KAP4 and CON21.
Stacked bar plots showing the numbers of genes undergoing promoter-type transitions between adjacent developmental stages in KAP4 (A) and CON21 (B)

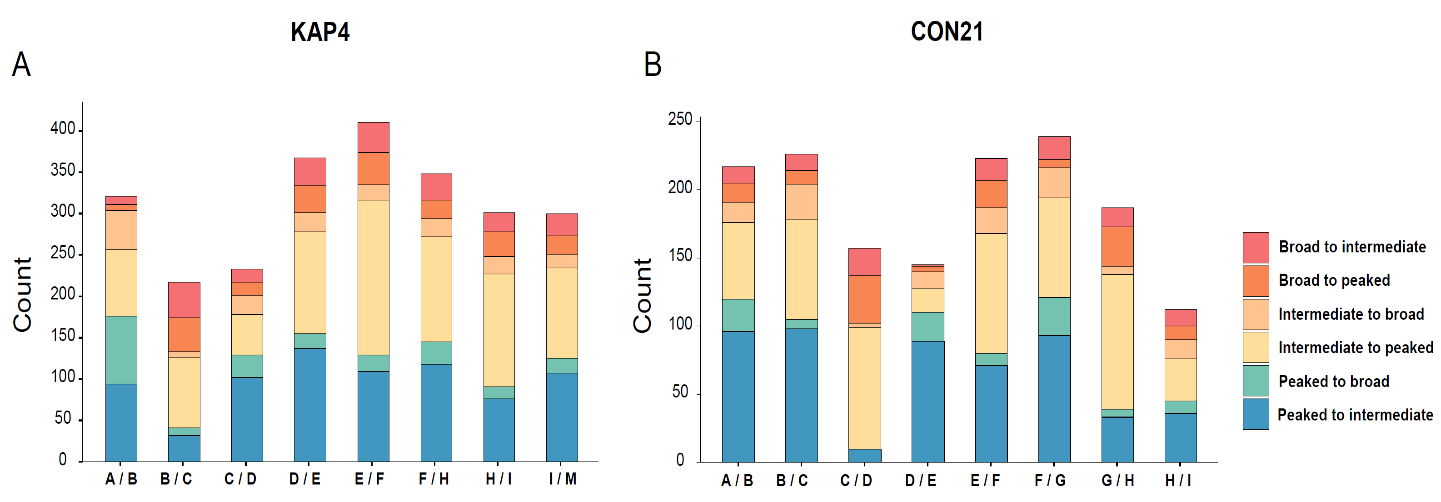

**Figure S9.** Expression divergence associated with conserved and divergent promoter types between KAP4 and CON21. Boxplots show the mean absolute expression difference, measured as |Δlog₂^(TPM + 1)^|, for genes expressed in both KAP4 and CON21 and grouped by promoter-type combination.

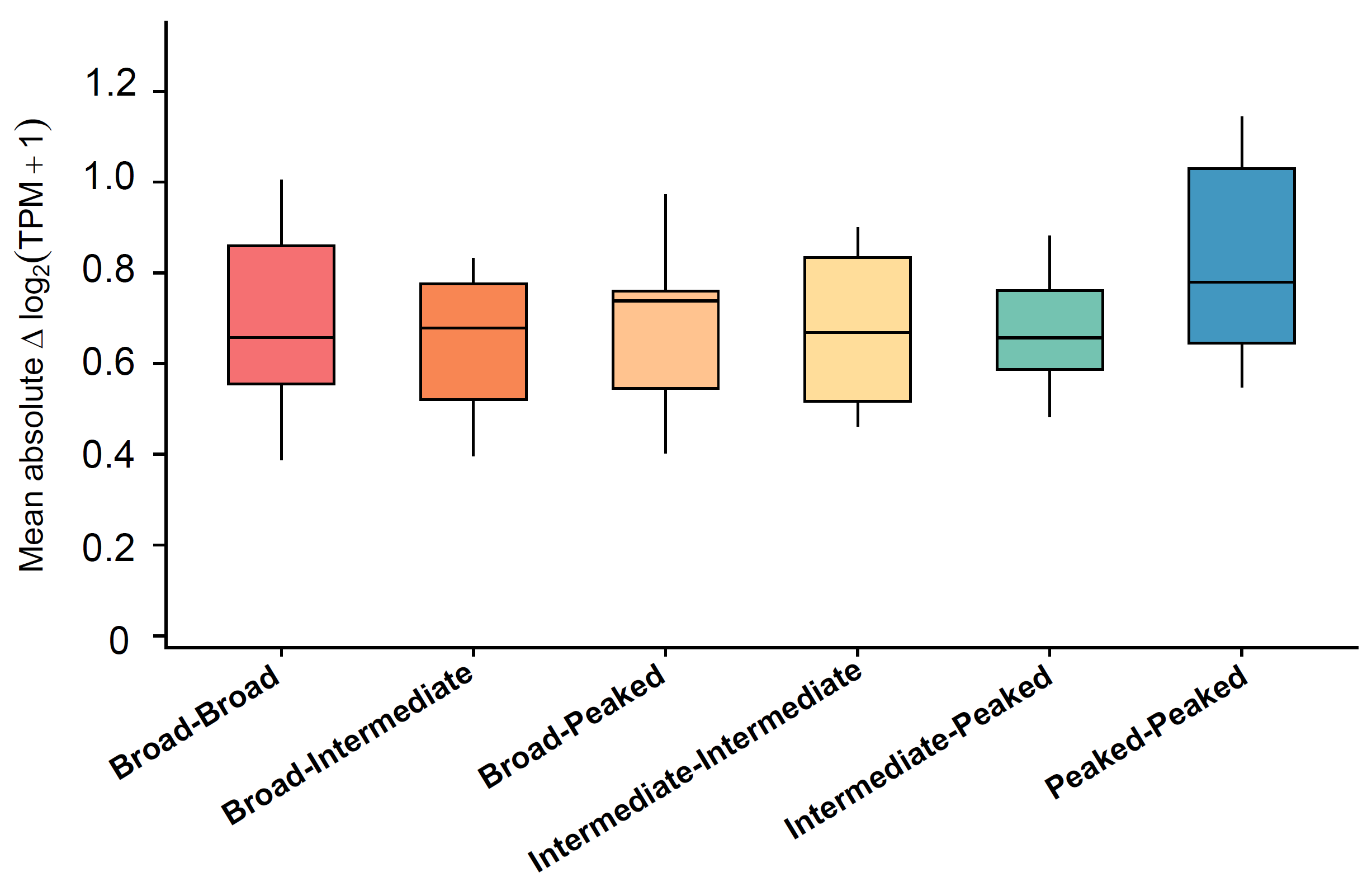
